# β-cell-specific *Ahr* expression is critical to high-fat diet-induced hyperinsulinemia

**DOI:** 10.64898/2026.06.25.734641

**Authors:** Ma. Enrica Angela Ching, Myriam P. Hoyeck, Lahari Basu, Jana Palaniyandi, Lili Grieco-St-Pierre, Renee Tejani, Erin van Zyl, Anastasiia Kostianets, Emilia Poleo-Giordani, Jennifer E. Bruin

## Abstract

**Objective:** The aryl hydrocarbon receptor (AhR) pathway primarily mediates pollutant responses by activating xenobiotic metabolism enzymes like cytochrome P450 1A1 and 1A2 (CYP1A). Although AhR has also been implicated in systemic metabolic dysfunction and is inducible in pancreatic islets, its role in islet physiology remains unclear.

**Methods:** We analyzed a publicly available bulk human islet transcriptomic dataset to identify pathways associated with *CYP1A1* expression. We also assessed islet responses to the pollutant 2,3,7,8-tetrachlorodibenzo-p-dioxin (TCDD) and glucolipotoxicity (GLT) *in vitro* using two mouse models: a global *Cyp1a1/1a2* double knockout (*Cyp*^KO^) model, which disrupts canonical AhR–CYP1A signaling in whole islets, and a β-cell-specific *Ahr* knockout (β*Ahr*^KO^) model, which abolishes AhR signaling selectively in β-cells. We then examined the role of β-cell *Ahr* in early adaptation to high-fat diet (HFD) feeding *in vivo*.

**Results:** Xenobiotic and nutrient metabolism pathways were enriched in donors with high *CYP1A1* expression. Global *Cyp1a1/1a2* deletion increased susceptibility of female mouse islets to TCDD-induced impairments in insulin secretion but had minimal effects on GLT responses in either sex. In contrast, β-cell *Ahr* deletion did not affect islet responses to TCDD, but exacerbated GLT-induced islet dysfunction in male islets and increased baseline insulin secretion in both vehicle- and GLT-exposed female islets *in vitro*. Lastly, β-cell *Ahr* deletion prevented adaptive HFD-induced hyperinsulinemia in both sexes *in vivo*.

**Conclusion:** Islet AhR signaling shapes responses to chemical and nutrient stressors in a context- and sex-dependent manner. While the canonical AhR–CYP1A axis supports female islet resilience to TCDD, β-cell AhR signaling more broadly regulates nutrient stress responses in both sexes.

## 1. INTRODUCTION

Type 2 diabetes (T2D) is a complex condition influenced by diverse intrinsic and extrinsic factors. For instance, genetic predisposition, consumption of energy-dense diets, and environmental pollutant exposure can all contribute to the onset and progression of T2D. One of the hallmarks of diabetes is impaired glucose regulation; glucose homeostasis is largely maintained by pancreatic islets, which are clusters of highly specialized hormone-producing cells [1,2]. β-cells secrete insulin in response to increased blood glucose levels and make up ~60-85% of islet cells in mice and humans [1–4]. Due to their high secretory demand, β-cells experience a substantial metabolic burden, rendering them highly susceptible to environmental stressors [5,6].

The aryl hydrocarbon receptor (AhR) pathway provides a potential mechanism through which environmental factors can influence islet health and function. The AhR is a ligand-activated transcription factor that binds to a diverse array of endogenous (e.g., tryptophan metabolites, eicosanoids) and exogenous (e.g., pharmaceuticals, environmental pollutants) compounds [7–10]. Although AhR is classically associated with xenobiotic metabolism and has been studied predominantly in the liver, recent evidence indicates AhR also regulates broader physiological processes, such as immune function and energy balance [9,11–13]. Importantly, the AhR pathway was shown to be inducible in both human and mouse islets [14,15], suggesting a direct role for AhR in islet biology.

AhR signaling can be broadly divided into canonical and non-canonical pathways. In the canonical AhR pathway, AhR is found in the cytosol bound to chaperone proteins [9,11–13]. Upon ligand binding, AhR translocates into the nucleus, dissociates from its chaperones, and forms a complex with aryl hydrocarbon receptor nuclear translocator (ARNT). The AhR–ARNT complex then binds to xenobiotic response elements (XREs) and recruits co-activators to initiate the transcription of target genes, such as xenobiotic metabolism enzymes, cytochrome P450 1A1 and 1A2 (*CYP1A*). In contrast, the non-canonical AhR pathway includes processes independent of AhR–ARNT–XRE binding and can act through genomic and non-genomic mechanisms. In the nucleus, AhR can interact with alternative transcriptional regulators, including members of the nuclear factor kappa B (NFκB) family and nuclear receptors, such as estrogen receptor alpha (ERα), to regulate inflammation, cell proliferation, and energy homeostasis [9,11–13]. AhR can also act through interactions with cytosolic factors. For instance, AhR activation can modulate Ca^2+^ channel activity and increase intracellular Ca^2+^ concentrations [9,16], which can impact β-cell insulin secretion.

The environmental pollutant 2,3,7,8-tetrachlorodibenzo-*p-*dioxin (TCDD) is the most potent inducer of the canonical AhR–CYP1A pathway, and exposure to TCDD and structurally-related chemicals has been associated with increased T2D incidence in humans [17–19]. In support of this, our lab has shown that 48 hours of TCDD exposure *in vitro* increases CYP1A1 activity in mouse and human islets, and this increase coincided with impaired glucose-stimulated insulin secretion (GSIS) [14]. Of note, while TCDD reliably activates CYP1A1 in islets, its effects on islet function and glucose regulation are less consistent. In another cohort of human donor islets, *in vitro* TCDD treatment induced *CYP1A1* without altering GSIS [20]. In mice, both chronic exposure to low-dose TCDD injections twice a week for 12 weeks and an acute single high-dose TCDD injection increased *Cyp1a1* expression in isolated islets. However, only the single-high dose injection led to impaired glucose homeostasis, reduced plasma insulin levels, and reduced GSIS in isolated islets [14,21]. Interestingly, these TCDD-induced changes in systemic glucose regulation and islet function were abolished when *Ahr* was deleted specifically in β-cells, suggesting that β-cell-specific AhR signaling mediates the effects of TCDD on islet function [21]. Altogether, these results demonstrate that the relationship between TCDD, AhR signaling, and islet function is complex and requires further elucidation.

Rodent studies demonstrate that AhR activity influences susceptibility to HFD-induced metabolic dysfunction [22–25]. Mice with a higher-affinity *Ahr* allele gained more weight and accumulated more hepatic fat on a HFD than mice with a lower-affinity *Ahr* allele [26], whereas administration of an AhR antagonist in HFD prevented obesity and reduced liver fat irrespective of allele affinity [23,27,28]. Likewise, global *Ahr* knockout mice exhibited less weight gain, reduced hepatic steatosis, and enhanced insulin sensitivity and glucose tolerance on HFD compared to wildtype controls [22,24]. Global knockout of downstream AhR targets, *Cyp1a1* and *Cyp1a2*, did not affect HFD-induced weight gain or hepatic steatosis but alleviated HFD-induced glucose intolerance in female mice [29]. Together, these findings suggest that AhR-driven metabolic phenotypes are not fully explained by the canonical AhR–CYP1A axis, emphasizing the need to further characterize how canonical and non-canonical AhR signaling contribute to glucose regulation.

Given the central role of islets in maintaining glucose homeostasis, we sought to understand how canonical and non-canonical AhR signaling influences islet and β-cell biology. We first analyzed a publicly available human islet transcriptomics dataset [30] to identify pathways associated with AhR activation, using *CYP1A1* expression as a biomarker of AhR activity. To further elucidate the role of AhR in islets, we used two complementary mouse models: a global *Cyp1a1/1a2* double knockout (*Cyp*^KO^) model to disrupt canonical AhR signaling through the AhR–CYP1A axis in islets, and a β-cell-specific *Ahr* knockout (β*Ahr*^KO^) model to assess the combined effects of canonical and non-canonical AhR pathways on β-cell function and physiology. Using these models, we examined how disruption of AhR signaling alters islet responses to chemical exposure and nutrient stress *in vitro*. We also assessed how β-cell-specific AhR signaling influences early metabolic adaptation to HFD feeding *in vivo*.

## 2. MATERIALS AND METHODS

### 2.1 Human islet transcriptomics

We used *CYP1A1* expression as a biomarker of canonical AhR pathway activation in human islets. Expression values were log_2_-transformed after adding a pseudocount of 1 (log_2_[*CYP1A1* + 1]) to accommodate zero values; for simplicity, we will refer to it as log_2_(*CYP1A1*) from here on.

We had access to two bulk human islet RNA sequencing (RNA-seq) datasets: one from the HumanIslets.com Consortium [31] and another from the Gene Expression Omnibus (GEO; accession: GSE50398) [30]. Although the HumanIslets.com dataset has a larger sample size (n = 371), *CYP1A1* was undetectable in 39.6% of samples and overall expression levels were lower than those observed in the GEO dataset (HumanIslets.com: mean = 4.5 ± 2.9, range: 0–11.0; GEO: mean = 9.3 ± 1.7, range: 4.6–13.1) **(Supplementary Figure 1)**. Therefore, we proceeded with the GEO dataset for subsequent analyses.

Individuals were binned into “low” and “high” *CYP1A1* expression groups based on log_2_(*CYP1A1*) expression. We evaluated two strategies for defining the threshold between these groups. First, we selected donors in the lowest 40% and highest 40% of the distribution (n = 35 per group), excluding the middle 20% (n = 18), to maximize contrast between donors with “high” versus “low” *CYP1A1* expression. Second, we used the 60^th^ percentile of log_2_(*CYP1A1*) expression as the cutoff and removed only the donors closest to this threshold (n = 4 removed; n = 84 retained) to maximize statistical power while minimizing exclusions. Since both approaches produced highly similar results **(Supplementary Figure 2; Figure 1C)**, we opted to use the 60th-percentile cutoff because it provides a clear separation between “low” and “high” *CYP1A1* groups while retaining the largest number of donors **(Figure 1Bi–ii)**. This approach resulted in a final cohort of 84 donors (33 females and 51 males). To assess sex differences, we also performed exploratory sex-stratified analyses **(Supplementary Figure 3)**.

**Figure 1.**
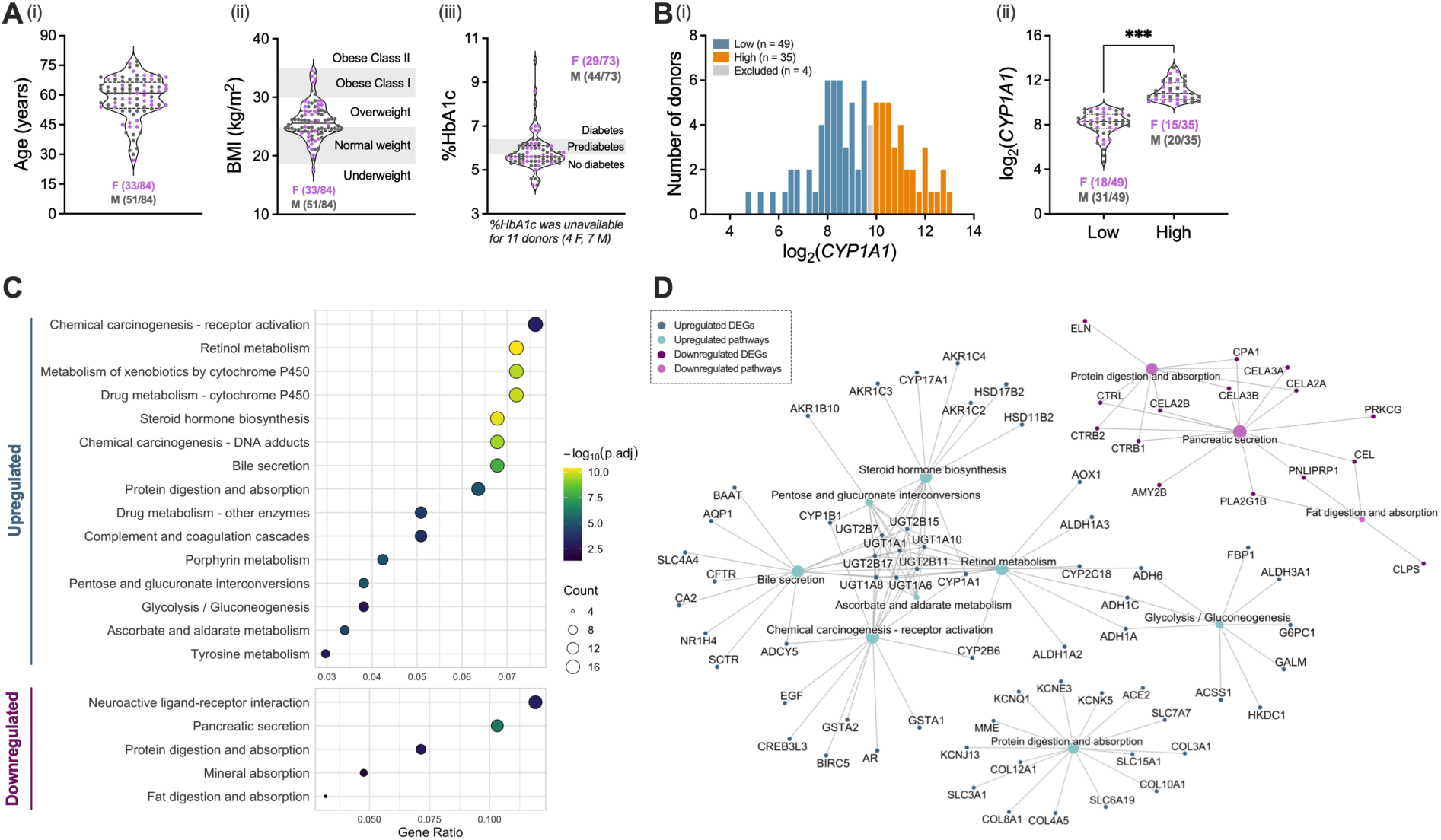
Human islet transcriptomics suggest *CYP1A1* expression is involved not only in xenobiotic metabolism but also in nutrient metabolism. Donor characteristics for 88 individuals (*Ai*–*iii*). Distribution of log_2_-transformed *CYP1A1* levels, log_2_(*CYP1A1*), showing 4 donors with intermediate *CYP1A1* levels (*Bi*). Comparison of log_2_(*CYP1A1*) between “low” and “high” *CYP1A1* groups after excluding intermediate donors (n = 84) (*Bii*). Dot plot of pathways enriched among upregulated differentially expressed genes (DEGs) (top) or downregulated DEGs (bottom), ordered by increasing GeneRatio (*C*). Category network plot showing DEGs associated with a subset of enriched pathways (*D*). Upregulated and downregulated networks were generated separately and combined for visualization; relative distances between networks are not meaningful. DEGs were identified using *DESeq2* (v1.50.2), pathway enrichment was performed using *clusterProfiler* (v4.18.4), and the graphs were generated with *enrichplot* (v1.30.5). All analyses were performed in R (v4.5.3) using RStudio (v2026.04.0+526).

Differential gene expression analysis was conducted using *DESeq2* (v1.50.2) using a model design controlling for age, sex, and BMI. Continuous variables (age, BMI) were z-transformed prior to modelling to reduce collinearity and improve model convergence. %HbA1c was excluded because it was moderately correlated with BMI (Pearson’s r = 0.49, p < 0.05) and missingness was unevenly distributed between the “low” (3/49 donors, 6%) and “high” (8/35 donors, 23%) *CYP1A1* groups. The threshold for differentially expressed genes (DEGs) was set at an FDR-adjusted p value (p_adj_) < 0.05 and an absolute log_2_ fold change (|log_2_FC|) > 0.58, corresponding to a ~1.5x linear fold change. We used *clusterProfiler* (v4.18.4) for pathway enrichment analyses and *enrichplot* (v1.30.5) for data visualization. All RNA-seq analyses were conducted in R (v4.5.3) using RStudio (v2026.04.0+526).

### 2.2 Animals

All mice were bred and housed in the Carleton University vivarium. The mice had *ad libitum* access to standard rodent chow (Harlan Laboratories, Teklad Diet, #2018, Madison, WI) and maintained on a 12-hour light/dark cycle. Cages were lined with corncob bedding (Inovativ, #7092, Azusa, CA), except for those used in the HFD study, which were lined with Tekfresh bedding (Inovativ, #7099P). All experiments were conducted in compliance with the standards established by Carleton University’s Animal Care Committee and the Canadian Council on Animal Care.

The *Cyp*^KO^ mouse line was kindly provided by Dr. Frank Gonzalez from the University of Cincinnati. This strain was originally generated by Dr. Daniel Nebert’s group using Cre-*lox*P technology on a C57BL/6J background [32]. The double-knockout genotype (*Cyp*^KO^, *Cyp1a1*^-/-^*1a2*^-/-^) was established by intercrossing *Cyp1a1*^-/-^ and *Cyp1a2*^-/-^ knockout mice, as described in Dragin et al. [32]. For all experiments, we used *Cyp*^WT^ (*Cyp1a1*^+/+^*1a2*^+/+^) littermate controls. Our colony has since undergone multiple backcrosses with C57BL/6N wild-type mice from Charles River Laboratories, resulting in a mixed C57BL/6J and C57BL/6N background.

The β*Ahr*^KO^ mouse model was previously validated in the Bruin Lab [21]. In this model, the *Ahr* gene is flanked by *lox*P sites in all cells, whereas *Cre recombinase* is under the control of the *Ins1* gene promoter. Because *Ins1* expression is restricted to β-cells, Cre-mediated *Ahr* deletion only occurs in β-cells. Both β*Ahr*^KO^ (*Ins1^Cre^*^/+^*Ahr*^fl/fl^) mice and β*Ahr*^WT^ (*Ins1^Cre^*^/+^*Ahr*^+/+^) mice have one copy of their *Ins1* gene replaced by *Cre recombinase*. As with the *Cyp*^KO^ colony, the background of these mice is also a mixture of C57BL/6J and C57BL/6N due to backcrossing.

For *in vitro* experiments, islets were isolated from multiple cohorts of female and male mice from both the *Cyp*^KO^ and β*Ahr*^KO^ colonies and exposed to either 2,3,7,8-tetrachlorodibenzo-*p-*dioxin (TCDD) or a high glucose, high fatty acid media to mimic glucolipotoxicity (GLT). TCDD was used as the model chemical as it is the most potent activator of the AhR pathway, while GLT was used to model an extreme nutrient-stress environment [33,34]. For TCDD experiments, islets were isolated from *Cyp*^WT^ and *Cyp*^KO^ mice aged 10–37 weeks and from β*Ahr*^WT^ and β*Ahr*^KO^ mice aged 18–24 weeks. For GLT experiments, islets were isolated from *Cyp*^WT^ and *Cyp*^KO^ mice aged 23–43 weeks and from β*Ahr*^WT^ and β*Ahr*^KO^ mice aged 13–27 weeks.

The β*Ahr*^KO^ model was also used in an *in vivo* HFD study. In brief, female and male β*Ahr*^KO^ mice and their β*Ahr*^WT^ littermate controls (8 to 16 weeks old) were fed a 10% low-fat diet (LFD) or a 60% HFD for 8 to 9 days. At endpoint, whole pancreas samples were collected for histology, and islets were isolated to assess islet function and gene expression. A detailed description of the HFD study is provided in Section 2.8.

### 2.3 Mouse islet isolation

Pancreatic duct injections were performed using collagenase (Sigma-Aldrich, #C7657) dissolved in Hanks’ balanced salt solution (HBSS: 137 mM NaCl, 5.4 mM KCl, 4.2 mM NaH_2_PO_4_, 4.1 mM KH_2_PO_4_, 10 mM HEPES, 1 mM MgCl_2_, 5 mM dextrose, pH 7.2) kept at 4°C. Following euthanasia by isoflurane overdose and cervical dislocation, the pancreas was inflated with the cold collagenase solution via the common bile duct, excised, then transferred to HBSS and kept on ice until further processing. Pancreas samples were incubated at 37°C to resume digestion for 10 min 30 sec, then vigorously shaken to further breakup tissue clumps. Cold HBSS with 1 mM CaCl_2_ was immediately added to prevent over-digestion. Digested pancreas tissues were then washed three times in cold HBSS with CaCl_2_ and resuspended in prewarmed (37°C) complete RPMI media (Wisent, #350-000-CL) supplemented with 1% Penicillin-Streptomycin (Gibco, #15-140-122) and 10% FBS (Sigma-Aldrich, #F1051). Next, histopaque (Sigma, #10771) was added to the tissue suspension, which was centrifuged at 2400 rpm for 18 min at 22°C to create a density gradient separating islets from exocrine tissue. The layer containing islets was collected, filtered through a 70-µm cell strainer, and resuspended in complete RPMI media. Islets were handpicked to >95% purity using dissecting microscopes (Zeiss Stemi 508), then incubated overnight at 37°C with 5% CO_2_ before performing *in vitro* experiments.

### 2.4 *In vitro* mouse islet culture and treatment conditions

For TCDD experiments, isolated mouse islets were transferred to 24-well non-TC (tissue culture) plates and cultured for 48 hours in complete RPMI media (Wisent, #350-000-CL) with 10 nM TCDD (310.6 µM TCDD in dimethyl sulfoxide [DMSO] stock, AccuStandard, #AD404SDMSO10X, Brockville, ON, Canada) or 0.03% (v/v) DMSO vehicle (Sigma-Aldrich, #276855). Islets were maintained at 37°C in 5% CO_2_, with a media change after ~24 hours. Following the 48-hour treatment, islets were washed three times with prewarmed Dulbecco’s Phosphate Buffered Saline (PBS; Sigma, #D8662) and handpicked for further experiments. For the *Cyp*^KO^ model, islets were assessed by perifusion (n = 7–10 mice per group; Section 2.5), microscopy-based PI-inclusion assays (n = 5–6 mice per group; Section 2.7A), and mitochondrial function assays (n = 10–14 mice per group; Section 2.6); for the β*Ahr*^KO^ model, islets were assessed by perifusion (n = 3–6 mice per group; Section 2.5; **Figure 2**).

**Figure 2.**
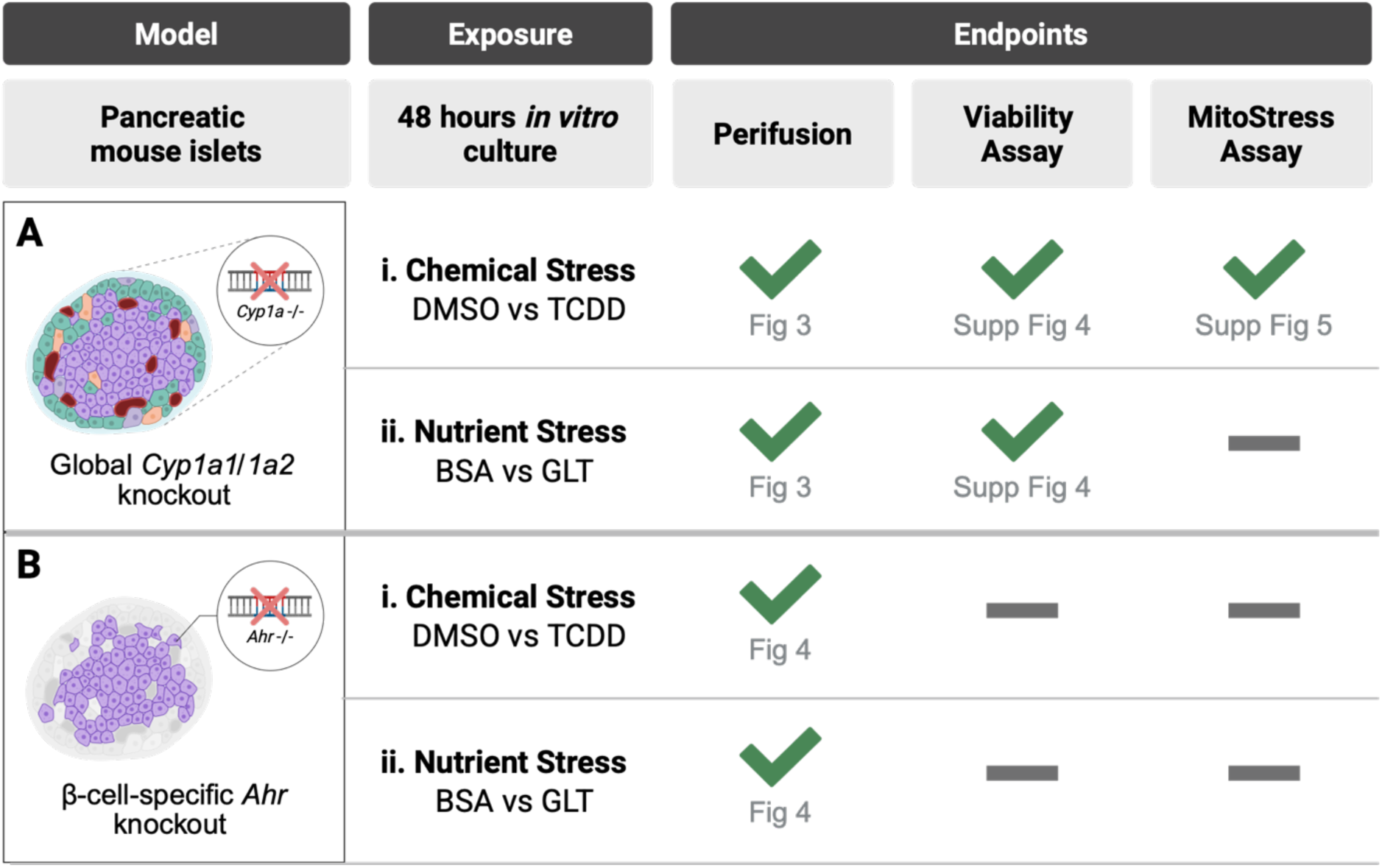
Schematic overview of *in vitro* experiments performed in isolated mouse islets from the global *Cyp1a1/1a2* knockout (*Cyp*^KO^) model and the β-cell-specific *Ahr* knockout (β*Ahr*^KO^) model. Islets were exposed to either a chemical or a nutrient stressor for 48 hours *in vitro*. Islet function was assessed by perifusion in all experiments (Ai–ii, Bi–ii). In *Cyp*^KO^ chemical stress experiments (Ai), islet cell viability was assessed with a microscopy-based propidium iodide (PI)-inclusion assay, and mitochondrial function was measured using a Seahorse XFe24 Analyzer. In *Cyp*^KO^ nutrient stress experiments (Aii), islet cell viability was quantified using a flow cytometry-based PI-inclusion assay.

For GLT experiments, islets were cultured in high-glucose, high-fatty acid media (41.1 mM glucose, 0.5 mM palmitate) or bovine serum albumin (BSA; Roche Diagnostics, #10775835001, Mannheim, Germany) vehicle control media (11.1 mM glucose, no fatty acid) for 48 hours. High glucose media was prepared by supplementing complete RPMI media (11.1 mM glucose) with D-(+)-glucose (2.5 M stock; Sigma-Aldrich, #G8769) to achieve a final concentration of 41.1 mM. Palmitate (Sigma-Aldrich, #P5585) was dissolved in 33 mM NaOH then complexed with 20% (w/v) BSA at a 5:1 molar ratio to make a 15.05 mM palmitate-BSA stock solution. To make the GLT media, the appropriate amount of the palmitate-BSA solution was added to the high glucose media to achieve a final concentration of 0.5 mM palmitate. For the BSA vehicle control, NaOH-BSA solution (33 mM NaOH in 20% BSA) was added to complete RPMI media. For the *Cyp*^KO^ model, islets were assessed by perifusion (n = 6 mice per group; Section 2.5) and flow-cytometry-based PI-inclusion assays (n = 4–5 mice per group; Section 2.7B); for the β*Ahr*^KO^ model, islets were assessed by perifusion (n = 3–5 mice per group; Section 2.5; **Figure 2**).

### 2.5 Glucose-stimulated insulin secretion perifusion assay

GSIS was dynamically assessed in isolated islets using a PERI4 automated perifusion system (Biorep Technologies). For each replicate, 70 islets were handpicked into microcentrifuge tubes and loaded into perifusion chambers between two layers of Bio-Gel P-4 polyacrylamide-based beads (Biorep Technologies, #PERI-BEADS) suspended in prewarmed Krebs-Ringer Bicarbonate HEPES (KRBB) buffer (0 mM glucose, 115 mM NaCl, 5 mM KCl, 2.5 mM CaCl_2_, 1 mM MgCl_2_, 10 mM HEPES, 24 mM NaHCO_3_, 0.1 % w/v BSA, pH 7.4). Islets were perifused with 2.8 mM low glucose (LG) KRBB for 40 minutes for equilibration, then for another 15 minutes to assess basal insulin secretion. The islets were then stimulated with 16.7 mM high glucose (HG) KRBB for a total of 45 minutes: 15 minutes to capture the 1^st^ phase and 30 minutes to capture the 2^nd^ phase of insulin secretion. After 25 minutes of recovery in LG KRBB, islets were depolarized with 30 mM KCl KRBB for 35 minutes, followed by a final 25-minute perifusion with LG KRBB. The flowrate was set to either 40 µL/min or 80 µL/min, depending on the assay step. Islets and perifusion solutions were maintained at 37°C and the collection plate was kept at 4°C throughout the experiment. Samples were stored at −80°C for long-term storage.

Insulin concentrations were measured either by rodent insulin chemiluminescence ELISA (ALPCO, #80-INSMR-CH10) or radioimmunoassay (Sigma-Aldrich, #RI-13K) according to manufacturer’s instructions.

### 2.6 Mitochondrial Function Assay

Mitochondrial function was assessed by measuring oxygen consumption rate (OCR) using a Seahorse XFe24 Analyzer (Agilent Technologies). For each replicate, 70 islets were handpicked into microcentrifuge tubes, then washed with prewarmed Seahorse XF RPMI medium (Agilent Technologies, #103576, pH 7.4) supplemented with 2 mM sodium pyruvate, 2 mM L-glutamine, 1% (v/v) FBS, and adjusted to a final glucose concentration of 2.8 mM using D-(+)-glucose (Sigma-Aldrich, #G8769). Islets were incubated at 37°C without CO_2_ for 1.5 hours in a 24-well islet capture plate (Agilent Technologies, #103518-100) coated with poly-D-lysine (Sigma-Aldrich, #P7280). After a media refresh, the plate was loaded into the Seahorse XFe24 Analyzer, and the protocol started with 5 cycles of basal OCR measurement. Islets were then exposed to sequential injections of 16.7 mM high glucose (HG, 6 cycles), 2.5 µM oligomycin (8 cycles), 3 µM carbonyl cyanide-4 (trifluoromethoxy) phenylhydrazone (FCCP, 5 cycles), and a combination of 3 µM rotenone and 3 µM antimycin A (Rot/AA, 6 cycles). The formulas used to calculate the mitochondrial function assay parameters are described in **Supplementary Table 2**.

### 2.7 Islet cell viability

#### 2.7A Microscopy-based PI-inclusion assay

Fifty islets per mouse per treatment were washed with prewarmed PBS (Sigma, #D8662) to remove residual media. To achieve a single cell suspension, islets were resuspended in 400 µL of Accutase (Corning, #25-058-CI) and incubated at 37°C for ~6 minutes, with gentle trituration every two minutes. Once ~70% of cells were dissociated, Accutase was neutralized with 600 µL of prewarmed complete RPMI media. Cells were pelleted at 2000 rpm for 1 min, washed with PBS, and resuspended in 250 µL media containing 0.5 µg/mL PI (Thermo Fisher, #P21493), 0.5 µM Hoechst (Thermo Fisher, #62249), and 1.25 µM Calcein AM (Thermo Fisher, #L3224A). The suspension was transferred to a 96-well non-TC-coated plate and incubated at room temperature for 1 hour in the dark before imaging. An Axio Observer 7 microscope (Carl Zeiss, Germany) was used to image 10% of each well. Live cells were identified as Hoechst⁺Calcein⁺PI⁻ and dead/dying cells as Hoechst⁺Calcein⁻PI⁺. Automated quantification of live and dead cells was performed using ZEN Blue 2.6 software (Carl Zeiss) and verified by manual counting by a single blinded observer. The proportion of dead/dying cells (% PI⁺) was calculated as: [# of dead cells / (# live cells + # dead cells)] × 100.

#### 2.7 B Flow cytometry-based PI-inclusion assay

Approximately 25–50 islets per mouse per treatment were cultured for 48 hours in the appropriate treatment or control media with 5 µg/mL PI (Thermo Fisher, #P21493). Three control conditions were included for each genotype: (1) a no PI negative control, (2) a PI-only control to measure baseline PI staining, and (3) a PI-positive oxidative stress control (250 µM H_2_O_2_ for 48 hours). Approximately 25–50 islets pooled from separate mice were used for each control condition. For the no PI and PI-only controls, the islets were cultured in untreated complete RPMI media, which did not contain the treatment (TCDD or GLT) or the vehicle (DMSO or BSA). The PI-positive oxidative stress control islets were cultured in the same untreated complete RPMI media but with the addition of 250 µM H_2_O_2_. All islets were dispersed into single cells using the same protocol as described in (A). The resulting single cell suspension was pelleted, resuspended in 200 µL PBS, and transferred to a 96-well plate for analysis using a BD Accuri C6 flow cytometer with a BD CSampler (BD Biosciences). Gating thresholds were defined using the controls and applied uniformly to all experimental samples to quantify the proportion of dead/dying cells (% PI⁺) per replicate. Background fluorescence was defined using the no-PI negative control, baseline PI fluorescence using the untreated PI-only control, and the PI-positive threshold using the H_2_O_2_-treated control.

### 2.8 *In vivo* HFD study using the β*Ahr*^KO^ model

Female and male β*Ahr*^KO^ and β*Ahr*^WT^ mice were transferred from a regular chow diet to a 10% LFD (Research Diets D12450J, New Brunswick, NJ) for an acclimatization period of 2 to 3 weeks. On day 0, a subset of mice was switched to a 60% HFD (Research Diets D12492, New Brunswick, NJ) for 8 to 9 days (see **Figure 5A** for study schematic). The diets differed in the relative proportion of fat and carbohydrate, but all other nutrient sources and proportions were matched. Mice in the LFD and HFD groups were matched based on average body weight and fasting blood glucose levels during the acclimatization period.

On day 8, a subset of mice was euthanized to collect whole pancreas for immunofluorescence histology (Section 2.10). Another subset of mice was euthanized on day 9 for pancreatic islet isolation (Section 2.3) for perifusion (Section 2.5) and quantitative real-time PCR (qPCR; Section 2.11).

### 2.9 *In vivo* metabolic assessments

Body weight and fasting blood glucose were measured following a 4-hour fast twice a week throughout the study. Saphenous blood glucose concentrations were measured with a StatStrip XPRESS handheld glucometer (Nova Biomedical, measuring range: 0.5–33.3 mmol/L). Whole-body fat composition was measured on days 0 and 6 using an EchoMRI-700 (EchoMRI LLC, Houston, TX). On day 7, mice were fasted for 6 hours prior to an intraperitoneal (ip) glucose tolerance test (GTT). Blood glucose levels were measured at 0, 15, 30, 60, and 90 min following a 2 g/kg ip glucose injection (DMVet Dextrose 50%, DIN: 02420880). Saphenous blood was obtained at all timepoints except 90 min to assess plasma insulin levels. Blood samples were collected using heparinized microhematocrit capillary tubes (Thermo Fisher, #22362566) and centrifuged (7000 rpm, 9 min, 4°C) to separate plasma. Plasma samples were stored at −20°C until insulin concentrations were measured by ELISA (ALPCO rodent ultrasensitive insulin ELISA, #80-INSMSU-E10, Salem, NH).

### 2.10 Histology

Pancreas tissues were fixed overnight in 4% paraformaldehyde (PFA; Fisher Scientific, #AAJ19943K2, Hampton, NA) at 4°C, then transferred to 70% ethanol until embedding. Tissues were processed and paraffin-embedded at the University of Ottawa Louise Pelletier Histology Core Facility. Immunostaining was performed as described previously [14] using a 10-min heat-induced epitope retrieval (HIER) step. To measure β-cell proliferation, HIER was performed in 1X TE buffer (10 mM Tris base, 1 mM EDTA tetrasodium, 0.05% [v/v] Tween 20, pH = 9), and slides were co-stained with rabbit anti-insulin (Ins; Cell Signaling Technology, C27C9, #3014S, 1:200 dilution, Danvers, MA) and mouse anti-proliferating cell nuclear antigen (PCNA; BD Transduction Laboratories, #610665, 1:100 dilution, Mississauga, ON). To measure average islet size and relative α- and β-cell area per islet, HIER was performed in 1X citrate buffer (10 mM citrate, 0.05% [v/v] Tween 20, pH = 6), and slides were co-stained with rabbit anti-insulin and mouse anti-glucagon (Gcg; Sigma-Aldrich, #G2654, 1:250 dilution). Following an overnight incubation in primary antibodies, sections were incubated for an hour in the secondary antibodies: goat anti-rabbit IgG (H+L) AF568 (Invitrogen, #A11011, 1:1000 dilution; Carlsbad, CA, USA) and goat anti-mouse IgG (H+L) AF488 dilution (Invitrogen, # A11029, 1:1000 dilution). Slides were mounted with VECTASHIELD HardSet Antifade Mounting Medium with DAPI (Vector Laboratories, #H-1500-10; Newark, CA, USA) and imaged using a ZEISS Axio Observer 7 (Carl Zeiss).

Images were manually quantified using ZEN Blue 2.6 software (Carl Zeiss). Islet cells with fluorescence above background levels were considered positive for their respective protein marker (Ins⁺, PCNA⁺, or Gcg⁺). Proliferating β-cells were defined as islet cells that were Ins⁺PCNA⁺, whereas non-proliferating β-cells were Ins⁺PCNA⁻. The proportion of proliferating cells per islet was calculated as: [# Ins⁺PCNA⁺ cells / (# Ins⁺PCNA⁺ + # Ins⁺PCNA⁻)] × 100. To quantify islet size and relative α- and β-cell proportions per islet, β-cells were defined as Ins⁺Gcg⁻ and α-cells as Ins⁻Gcg⁺. The proportion of hormone⁺ (i.e., Ins⁺ or Gcg⁺) area was calculated as: (hormone⁺ area / total islet area) × 100.

### 2.11 Quantitative real-time PCR

Islets for qPCR were lysed in buffer RLT (RNeasy Micro Kit, QIAGEN, #74004; Hilden, Germany) supplemented with 1% β-mercaptoethanol (Sigma Aldrich, #M3148, St. Louis, MO) and stored at −80°C. Total RNA was extracted using the RNAeasy Micro Kit (QIAGEN, #74004), and cDNA was synthesized using the iScript gDNA Clear cDNA Synthesis Kit with DNase treatment (Bio-Rad, #1725035, Hercules, CA, USA), following the manufacturers’ instructions. Relative mRNA levels were quantified using the SsoAdvanced Universal SYBR Green Supermix (Bio-Rad, #1725271) with Precision Blue Real-Time PCR Dye (Bio-Rad, #1725275) on a CFX384 real-time qPCR detection system (Bio-Rad, #43174). Gene expression was analyzed using the 2^-ΔΔCt^ method, with *Ppia* as the reference gene. Primers were synthesized by Integrated DNA Technologies (IDT; Coralville, IA, USA); primer sequences are in **Supplementary Table 3**.

### 2.12 Statistical analyses

Statistical analyses were performed using GraphPad Prism (version 10.6.1 for macOS) or RStudio (2025.09.1+401; R version 4.4.2). Treatment and genotype effects were assessed by repeated-measures two-way ANOVA or a mixed-effects model, followed by an uncorrected Fisher’s least significant difference (LSD) test in Prism. Analyses involving three factors (genotype, treatment, and time) were performed in RStudio using linear mixed-effects models, followed by Tukey’s post hoc tests on estimated marginal means for pairwise comparisons. Statistical significance was set at *p* < 0.05. Data are presented as mean ± SEM; individual datapoints represent biological replicates.

## 3. RESULTS

### 3.1 *CYP1A1* expression in human islets is associated with pathways related to both xenobiotic and nutrient metabolism

The human islet donor population (n = 84) had a mean age of 58 ± 11 years (range: 27–77 years), a mean body mass index (BMI) of 26.0 ± 3.2 kg/m² (range: 17.6–34.6 kg/m²), and a mean percent hemoglobin A1c (HbA1c) level of 5.8 ± 0.9% (range: 4.3–10.0%) **(Figure 1A)**. Based on BMI classifications set by the World Health Organization (WHO) [35], 1.2% of donors (n = 1) had a BMI in the underweight range (< 18.5 kg/m²), 44.0% (n = 37) of donors were in the normal weight range (18.5–24.9 kg/m²), 46.4% (n = 39) were in the overweight range (25–29.9 kg/m²), and 8.3% (n = 7) were in the class I obesity range (30–34.9 kg/m²) **(Figure 1Aii)**. According to guidelines from the American Diabetes Association (ADA) [36], 44.0% (n = 37) of donors had HbA1c values in the non-diabetes range (< 5.7%), 32.1% (n = 27) were in the prediabetes range (5.7–6.4%), and 10.7% (n = 9) were in the diabetes range (HbA1c ≥ 6.5%). Glycemic status was unknown for 13.1% (n = 11) of donors **(Figure 1Aiii)**.

To investigate whether endogenous AhR pathway activity influences gene expression profiles in human islets, we divided donors into “low *CYP1A1*” and “high *CYP1A1*” groups based on log_2_(*CYP1A1*) values **(Figure 1Bi)**. The “low *CYP1A1*” group had a mean log_2_(*CYP1A1*) expression of 8.1 ± 1.1 (range: 4.6–9.5), whereas the “high *CYP1A1*” group had a mean of 11.0 ± 0.9 (range: 9.9–13.1) **(Figure 1Bii)**. We identified 739 DEGs between human islets with low versus high *CYP1A1* levels (p_adj_ < 0.05 and |log_2_FC| > 0.58), of which 361 (48.9%) were successfully mapped to KEGG pathways. The top 50 mapped DEGs based on absolute log_2_FC are listed in **Supplementary Table 1**.

Although the most strongly enriched pathways were related to xenobiotic metabolism and chemical carcinogenesis, several nutrient metabolism pathways were also enriched in donors with high versus low *CYP1A1* expression. Among DEGs upregulated in high *CYP1A1* donors relative to low *CYP1A1* donors, enriched pathways included retinol metabolism, protein digestion and absorption, pentose and glucuronate interconversions, glycolysis/gluconeogenesis, and ascorbate and aldarate metabolism. In contrast, pathways enriched among DEGs downregulated in donors with high versus low *CYP1A1* expression were primarily related to nutrient metabolism, including pancreatic secretion, protein digestion and absorption, mineral absorption, and fat digestion and absorption **(Figure 1C)**.

Exploratory sex-stratified pathway enrichment analyses revealed several core pathways that were impacted in both sexes but also showed sex-specific patterns **(Supplementary Figure 2A–B)**. Pathways enriched in the pooled analysis and in both sexes individually included xenobiotic metabolism pathways and retinol metabolism. In females, the top enriched pathways among DEGs upregulated in the high-*CYP1A1* group were steroid hormone biosynthesis and retinol metabolism, while female-specific pathways were mostly related to hormone and nutrient metabolism (thyroid hormone synthesis, pyruvate metabolism, and fatty acid degradation) **(Supplementary Figure 2A)**. In males, the most highly enriched pathways donors with high versus low *CYP1A1* expression were related to xenobiotic metabolism, and male-specific pathways were broadly associated with the immune system, extracellular matrix (ECM), and heme-related metabolism (e.g., integrin signaling, ECM-receptor interaction, *Staphylococcus aureus* infection, complement and coagulation cascades, and porphyrin metabolism) **(Supplementary Figure 2B)**.

We also generated network plots using a subset of enriched pathways from the pooled analysis to visualize the relationships between these pathways and their associated DEGs **(Figure 1D)**. Xenobiotic metabolism pathways shared several genes with nutrient metabolism pathways, suggesting overlap between detoxification pathways and metabolic processes in human islets. For instance, several phase II xenobiotic metabolism enzymes, including *UGT1A8*, *UGT1A10*, and *UGT2B15*, appeared in retinol metabolism, pentose and glucuronate interconversions, ascorbate and aldarate metabolism, and bile secretion. Additionally, alcohol dehydrogenases, *ADH1A*, *ADH1C*, and *ADH6* were also present in the glycolysis/gluconeogenesis pathway **(Figure 1D)**. Finally, several DEGs were mapped exclusively to nutrient pathways, further suggesting that AhR activation in human islets not only induces xenobiotic detoxification processes but may also extend to physiological processes critical for maintaining islet homeostasis. To explore this further, we assessed how isolated islets from two different transgenic mouse models (*Cyp*^KO^, β*Ahr*^KO^) respond to either xenobiotic stress (TCDD) or nutrient stress (GLT) *in vitro* (**Figure 2**).

### 3.2 The AhR–CYP1A axis mediates TCDD-induced β-cell dysfunction in female mouse islets

To examine the role of CYP1A enzymes in whole islets under xenobiotic stress, we treated female and male *Cyp*^WT^ and *Cyp*^KO^ mouse islets *in vitro* for 48 hours with either DMSO or TCDD, the most potent activator of canonical AhR signaling **(Figure 2Ai)**. First, we confirmed that the TCDD treatment protocol did not affect islet cell viability, regardless of sex or genotype **(Supplementary Figure 4A–B)**. We then assessed islet function and found that in female islets, TCDD exposure led to modest reductions in insulin secretion, but the effect was more pronounced in *Cyp*^KO^ islets. In *Cyp*^WT^ islets, TCDD reduced only 1^st^ phase GSIS (t = 12–14 min; **Figure 3Ai, iii**), whereas in *Cyp*^KO^ islets, TCDD modestly suppressed insulin secretion during both the 1^st^ and 2^nd^ phases of GSIS (t = 14–36 min; **Figure 3Ai–iv**) and KCl-stimulated insulin secretion (t = 72–84 min; **Figure 3Ai, v**). In contrast, TCDD had no effect on insulin secretion in male *Cyp*^WT^ or *Cyp*^KO^ islets **(Figure 3B)**. These data suggest that the absence of *Cyp1a1*/*1a2* in female islets exacerbates TCDD-induced effects on GSIS.

**Figure 3.**
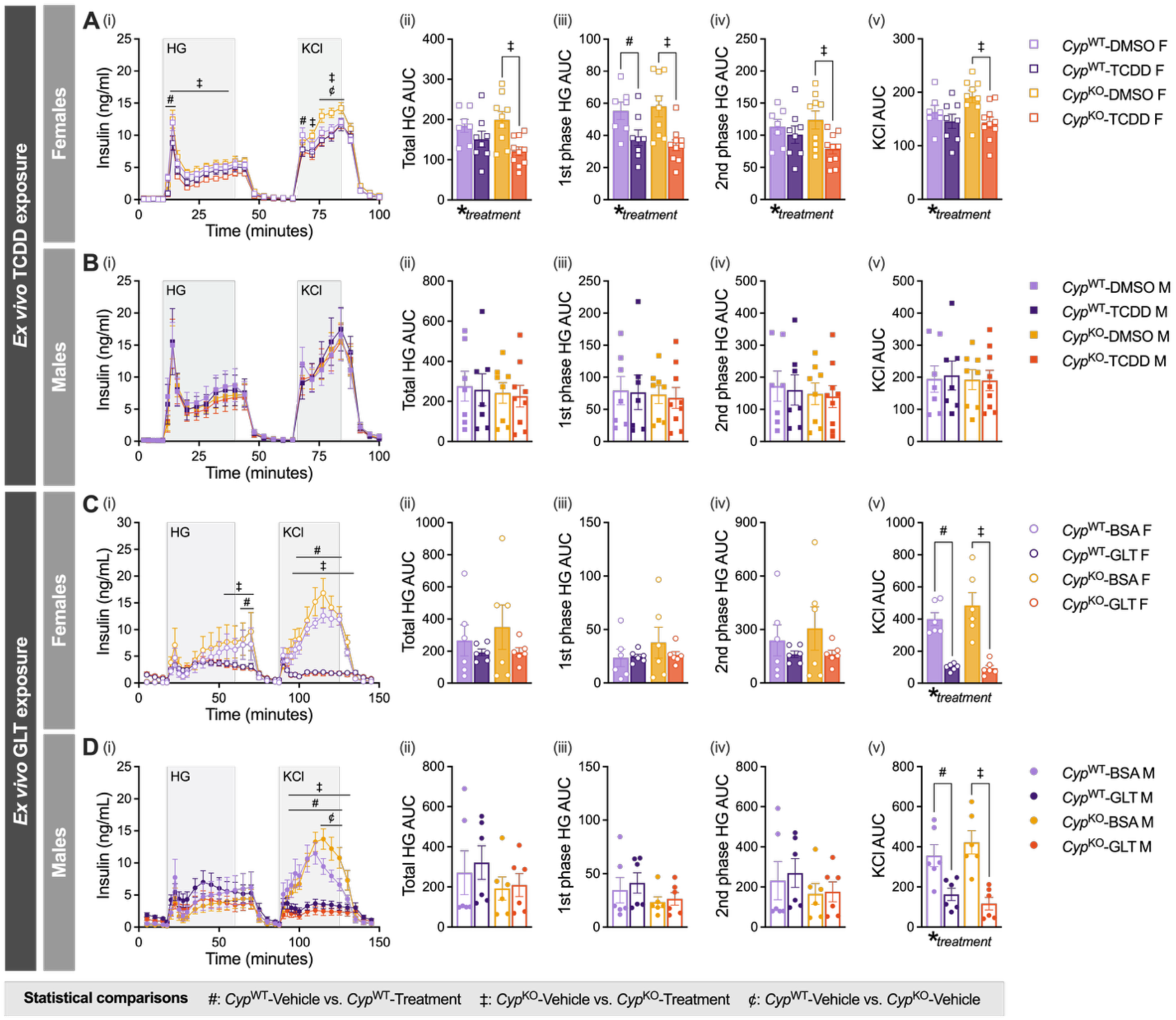
The AhR–CYP1A axis mediates TCDD-induced β-cell dysfunction in female mouse islets but does not influence islet responses to glucolipotoxicity (GLT) in either sex. β-cell function was assessed in *Cyp*^WT^ and *Cyp*^KO^ female (*A*, *C*) and male (*B*, *D*) mouse islets following 48 hours of *ex vivo* exposure to 2,3,7,8-tetrachlorodibenzo-*p-*dioxin (TCDD; *n*_females_ = 7–10 per group, *n*_males_ = 7–9 per group; *A*–*B*) or GLT (*n*_females_ = 6 per group, *n*_males_ = 6 group; *C*–*D*). Insulin concentration (ng/mL) (*i*) was measured every 2.5 to 5 minutes as islets were exposed to 2.8 mM low glucose, 16.7 mM high glucose (HG), or 30 mM KCl solutions. Area under the curve (AUC) values were calculated for total HG (t = 10–48 minutes; *Aii*, *Bii*), 1^st^ phase HG (t = 10–20 minutes; *Aiii*, *Biii*), 2^nd^ phase HG (t = 24–48 minutes; *Aiv*, *Biv*), and KCl-stimulated insulin secretion (t = 68–84 minutes; *Av*, *Bv*) in TCDD experiments, and for total HG (t = 17.5–80 minutes; *Cii*, *Dii*), 1^st^ phase HG (t = 17.5–27.5 minutes; *Ciii*, *Diii*), 2^nd^ phase HG (t = 30–85 minutes; *Civ*, *D*iv), and KCl-stimulated insulin secretion (t = 87.5–140 minutes; *Cv*, *Dv*) in GLT experiments. All data are presented as means ± SEM. Bar graphs show AUC values, with individual data points representing biological replicates (*ii*–*v*). The following statistical tests were used — line graphs (*i*): linear mixed-effects model followed by Tukey’s post hoc tests on estimated marginal means for pairwise comparisons at each timepoint. Bar graphs (*ii*–*v*): mixed effects model with uncorrected Fisher’s LSD. Statistically significant results (*p* < 0.05) are denoted by an asterisk (*) for overall two-way ANOVA effects or by the following symbols for pairwise comparisons — #*Cyp*^WT^-DMSO vs. *Cyp*^WT^-TCDD; ‡*Cyp*^KO^-DMSO vs. *Cyp*^KO^-TCDD; ¢*Cyp*^WT^-DMSO vs. *Cyp*^KO^-DMSO.

To investigate potential mechanisms underlying differences in insulin secretion between DMSO and TCDD-treated islets, we assessed mitochondrial function. Non-mitochondrial respiration, basal respiration, acute HG respiration, and ATP-linked respiration were similar in female islets between genotypes and treatments **(Supplementary Figure 5Ai–ii, v)**. TCDD-treated *Cyp*^WT^ female islets showed a ~1.25-fold increase in maximal OCR and a trending ~1.24-fold increase in spare respiratory capacity compared to their DMSO-treated counterparts, and this response to TCDD was absent in *Cyp*^KO^ female islets **(Supplementary Figure 5Avi–vii)**. In line with the perifusion data, mitochondrial function in male islets was not affected by genotype or treatment **(Supplementary Figure 5Bi–vii)**. Overall, our data suggest CYP1A1 enzymes are involved in supporting mitochondrial adaptation to xenobiotic stress in female mouse islets, which may contribute to the observed differences in islet function.

### 3.3 The AhR–CYP1A axis does not influence islet responses to *in vitro* GLT stress in either sex

Next, we assessed the role of CYP1A enzymes in islet responses to nutrient stress, by treating islets from female and male *Cyp*^WT^ and *Cyp*^KO^ mice *in vitro* for 48 hours with either BSA or GLT media (**Figure 2Aii).** We began by verifying that there were no significant differences in islet cell viability between vehicle- and GLT-treated islets regardless of sex or genotype **(Supplementary Figure 4C–D)**. In females, GLT treatment modestly reduced 2^nd^ phase insulin secretion compared to vehicle in both *Cyp*^WT^ and *Cyp*^KO^ **(Figure 3Ci)**, although this did not amount to an overall change in AUC **(Figure 3Ci, iv)**. The most pronounced effect of GLT was a marked suppression of KCl-stimulated insulin secretion irrespective of sex or genotype (KCl AUC reduced by ~75–80%; **Figure 3C–D, i, v**). The absence of genotype-driven effects suggests that the AhR–CYP1A axis does not play a major role in the response of mouse islets to GLT stress.

Thus far, our results demonstrate that the AhR–CYP1A axis in islets is involved in responding to xenobiotic stress in females, but had no effect on the GLT stress response in either sex, at least under the conditions tested in this experimental model. However, *Cyp1a1/1a2* represents only one downstream component of the AhR signaling pathway. To examine the broader role of AhR beyond CYP1A-mediated responses and more directly assess its role in β-cell physiology, we next repeated the *in vitro* TCDD and GLT experiments using the β*Ahr*^KO^ model.

### 3.4 β-cell-specific *Ahr* deletion does not alter islet responses to TCDD in either sex

First, we exposed female and male β*Ahr*^WT^ and β*Ahr*^KO^ islets to DMSO or 10 nM TCDD for 48 hours *in vitro* **(Figure 2Bi)**. TCDD exposure did not affect insulin secretion relative to DMSO controls in either sex or genotype **(Figure 4A–B)**. Interestingly, there was a trending overall genotype effect in females, with β*Ahr*^KO^ islets showing an increase in total HG AUC and 2^nd^ phase HG AUC relative to β*Ahr*^WT^ islets **(Figure 4Aii–iv)**. The KCl response was comparable between β*Ahr*^WT^ and β*Ahr*^KO^ female islets **(Figure 4Av)**. In male islets, neither genotype nor TCDD treatment affected insulin secretion **(Figure 4Bi–v)**. These results suggest that β-cell-specific *Ahr* deletion enhances GSIS in female islets under basal conditions but does not impact the overall islet response to TCDD in either sex.

**Figure 4.**
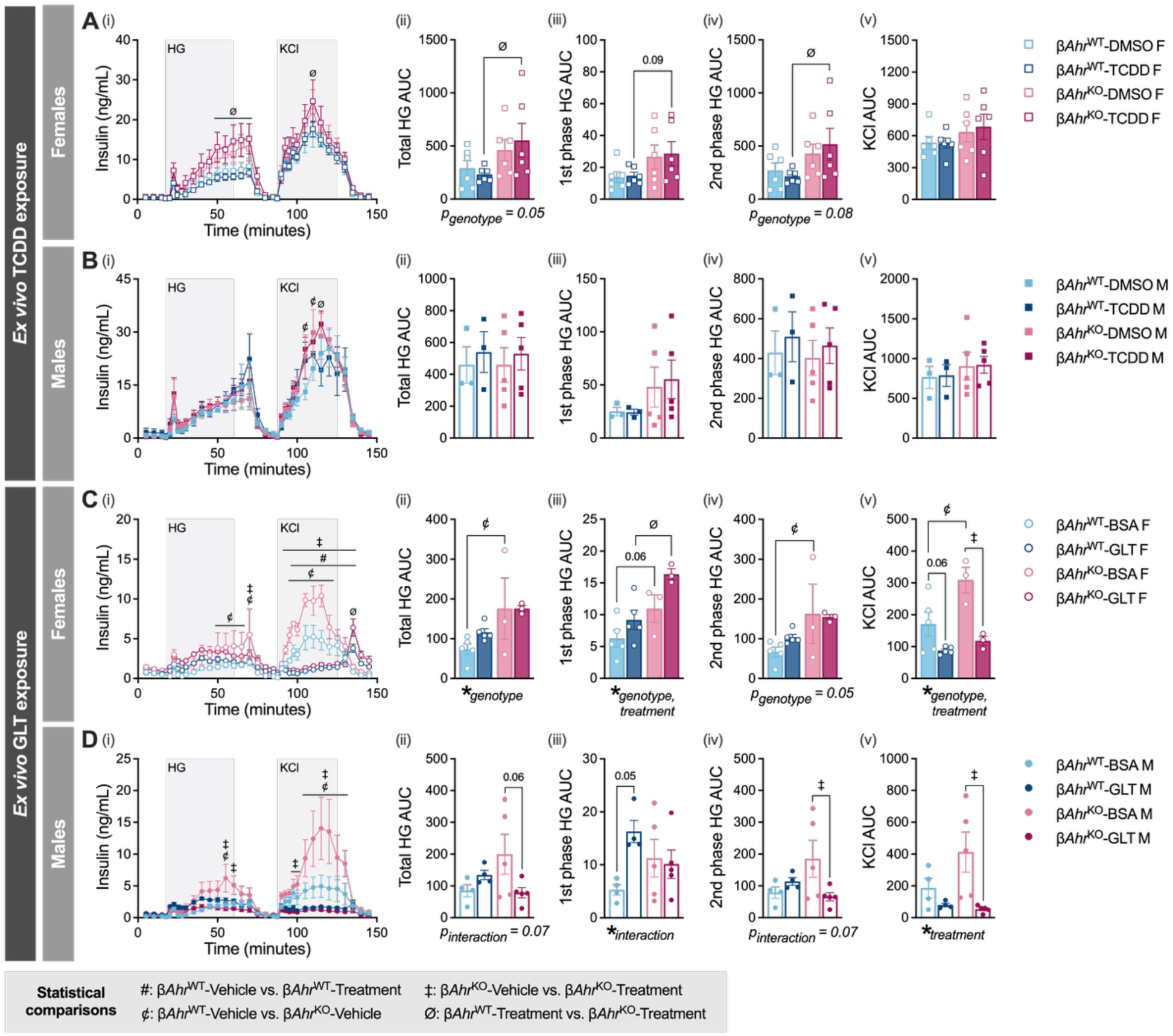
β-cell-specific *Ahr* contributes to islet responses to glucolipotoxicity *in vitro*. β-cell function was assessed in β*Ahr*^WT^ and β*Ahr*^KO^ female (*A*, *C*) and male (*B*, *D*) mouse islets following 48 hours of *ex vivo* exposure to 2,3,7,8-tetrachlorodibenzo-*p-*dioxin (TCDD; *n*_females_ = 6 per group, *n*_males_ = 3–5 per group; *A*–*B*) or GLT (*n*_females_ = 3–5 per group, *n*_males_ = 4–5 group; *C*–*D*). The vehicle controls were dimethyl sulfoxide (DMSO) for TCDD and bovine serum albumin (BSA) for GLT. Insulin concentration (ng/mL) (*i*) was measured every 2.5 to 5 minutes as islets were exposed to 2.8 mM low glucose, 16.7 mM high glucose (HG), or 30 mM KCl solutions. Area under the curve (AUC) values were calculated for total HG insulin secretion (t = 17.5–80 minutes) (*ii*), 1^st^ phase HG insulin secretion (t = 17.5–27.5 minutes) (*iii*), 2^nd^ phase HG insulin secretion (t = 30–85 minutes) (*iv*), and KCl insulin secretion (t = 87.5–140 minutes) (*v*). All data are presented as means ± SEM. Bar graphs show AUC values, with individual data points representing biological replicates (*ii*–*v*). The following statistical tests were used — line graphs (*i*): linear mixed-effects model followed by Tukey’s post hoc tests on estimated marginal means for pairwise comparisons at each timepoint. Bar graphs (*ii*–*v*): repeated measures two-way ANOVA with uncorrected Fisher’s LSD. Statistically significant results (*p* < 0.05) are denoted by an asterisk (*) for overall two-way ANOVA effects or by the following symbols for pairwise comparisons — #β*Ahr*^WT^-Vehicle vs. β*Ahr*^WT^-Treatment; ‡β*Ahr*^KO^-Vehicle vs. β*Ahr*^KO^-Treatment; ¢β*Ahr*^WT^-Vehicle vs. β*Ahr*^KO^-Vehicle; Øβ*Ahr*^WT^-Treatment vs. β*Ahr*^KO^-Treatment.

### 3.5 β-cell-specific *Ahr* contributes to islet responses to GLT *in vitro*

We next assessed whether the AhR pathway in β-cells mediates the response of islets to nutrient stress. Female and male β*Ahr*^WT^ and β*Ahr*^KO^ islets were exposed to either BSA or GLT media for 48 hours *in vitro* **(Figure 2Bii)**. Consistent with the TCDD experiments **(Figure 4Aii, iv)**, β*Ahr*^KO^ female islets showed increased GSIS compared to β*Ahr*^WT^ islets, with a significant overall genotype effect on total, 1^st^ phase, and 2^nd^ phase HG AUCs **(Figure 4Ci–iv)**. In female islets, GLT treatment did not alter GSIS in either genotype but caused a drastic ~50-60% impairment in KCl-stimulated insulin secretion in both β*Ahr*^WT^ and β*Ahr*^KO^ islets relative to vehicle **(Figure 4Ci, v)**. In male islets, there were no significant differences in insulin secretion between β*Ahr*^WT^ and β*Ahr*^KO^ islets under BSA control conditions **(Figure 4Di–vi)**. Following GLT exposure, β*Ahr*^WT^ male islets exhibited a ~3-fold increase in 1^st^ phase HG AUC **(Figure 4Diii)** compared to BSA-exposed β*Ahr*^WT^ islets, with no changes in total or 2^nd^ phase HG AUCs **(Figure 4Dii, iv)**. In contrast, GLT did not affect 1^st^ phase GSIS in β*Ahr*^KO^ islets **(Figure 4Diii)** but rather caused a 60% decrease in total and 2^nd^ phase HG AUC relative to BSA-treated β*Ahr*^KO^ islets **(Figure 4Dii, iv)**. GLT also reduced KCl-stimulated insulin secretion in both genotypes, although the magnitude of the suppression was greater in β*Ahr*^KO^ islets **(Figure 4Dv)**. Overall, these results suggest β-cell AhR signaling contributes to basal insulin secretion in females, while regulating islet responses to *in vitro* nutrient stress in males.

### 3.6 β-cell-specific *Ahr* deletion did not influence HFD-induced increases in body weight, fat mass, or fasting glycemia

To further investigate the role of β-cell AhR signaling in early metabolic adaptation to nutrient stress, we performed a ~1-week HFD study *in vivo* using the β*Ahr*^KO^ model. Female and male β*Ahr*^WT^ and β*Ahr*^KO^ mice were transitioned from chow to 10% LFD for at least 7 days before a subset of mice were transferred to 60% HFD for 8 to 9 days (*n*_females_ = 8–9 per group, *n*_males_ = 7–10 per group; **Figure 5A**). Both female and male mice showed increased body weight and % fat mass after 7 days of HFD feeding regardless of genotype **(Figure 5B–C, E–F)**. Only HFD-fed males exhibited fasting hyperglycemia on day 7 **(Figure 5Dii, Gii)**, demonstrating that metabolic dysfunction is accelerated in males compared to females.

**Figure 5.**
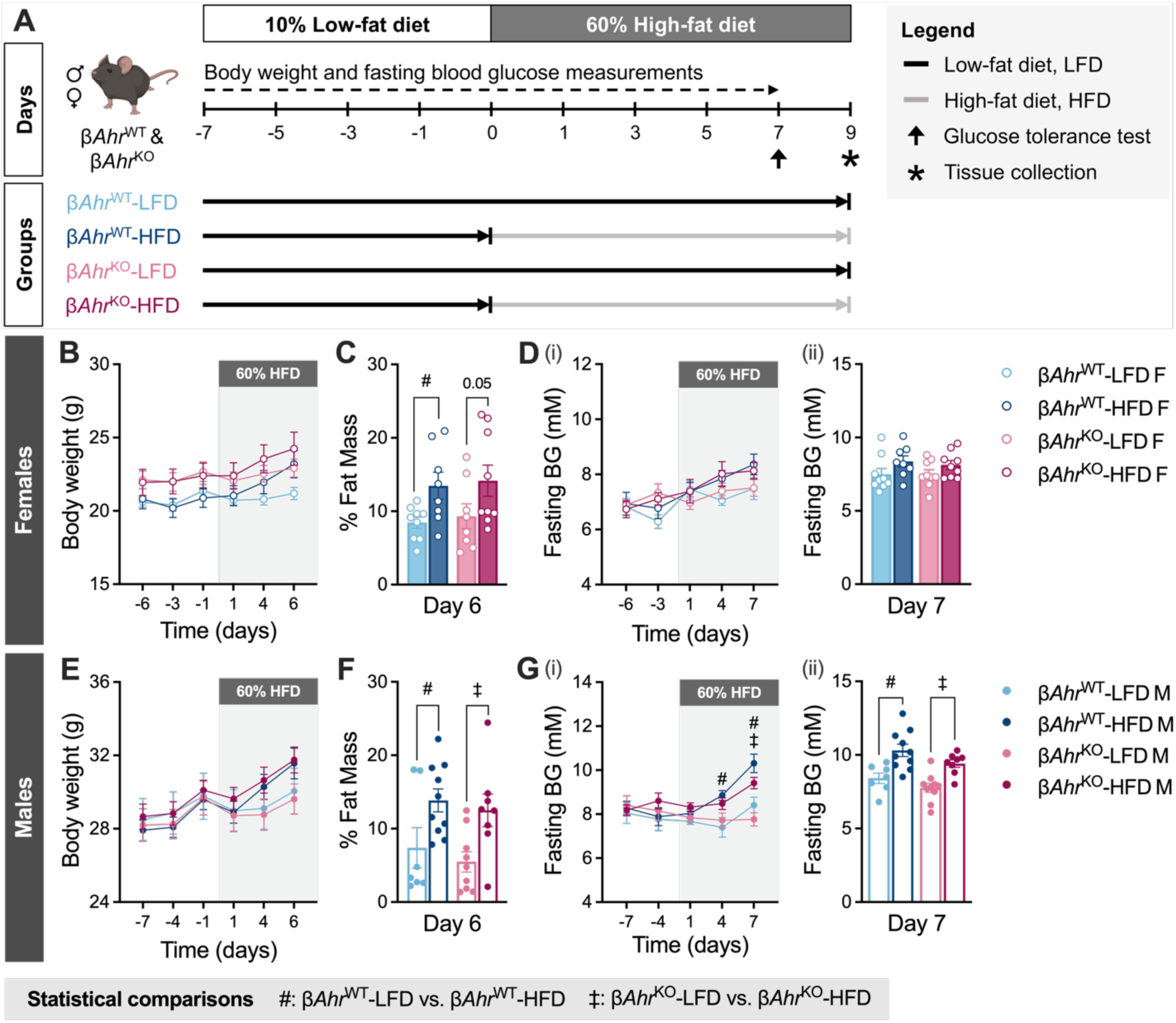
β-cell-specific *Ahr* deletion did not influence HFD-induced increases in body weight, fat mass, or fasting glycemia. Male and female β*Ahr*^WT^ and β*Ahr*^KO^ mice were fed a low-fat diet (LFD) for at least a week before a subset of mice were transferred to high-fat diet (HFD) for 7 days (*n*_females_ = 8–9 per group, *n*_males_ = 7–10 per group). Mice were euthanized for tissue collection on *day 9* (*A*). Body weight tracking (*B*, *E*) and % fat mass on *day 6* (*C*, *F*). Fasting blood glucose (BG) measurements (*D*, G) throughout the study (*i*) and on *day 7* (*ii*). All data are presented as means ± SEM. Individual data points represent biological replicates. The following statistical tests were used — line graphs (*B*, *Di*, *E*, *Gi*): mixed-effects model with Tukey’s post hoc test. Bar graphs (*C*, *Dii*, *F*, *Gii*): ordinary two-way ANOVA with uncorrected Fisher’s LSD. Statistically significant results (*p* < 0.05) are denoted by the following symbols for pairwise comparisons — #β*Ahr*^WT^-LFD vs. β*Ahr*^WT^-HFD; ‡β*Ahr*^KO^-LFD vs. β*Ahr*^KO^-HFD.

### 3.7 Deleting *Ahr* in β-cells prevented HFD-induced hyperinsulinemia in both sexes *in vivo*, but had sex-specific effects on insulin secretion *ex vivo*

On day 7, HFD-fed female and male β*Ahr*^WT^ and β*Ahr*^KO^ mice were glucose intolerant compared to their LFD-fed counterparts **(Figure 6A, D)**. HFD feeding increased the overall glucose excursion by ~1.45-fold in females and ~1.6-fold in males. Although the effect of HFD on glucose tolerance was comparable between β*Ahr*^WT^ and β*Ahr*^KO^ mice, the accompanying plasma insulin response showed clear genotype-based differences, with β*Ahr*^KO^ mice lacking the hyperinsulinemia observed in β*Ahr*^WT^ mice. In females, HFD feeding modestly elevated plasma insulin at 60 min post-glucose in β*Ahr*^WT^, but not β*Ahr*^KO^ mice **(Figure 6B)**. This phenotype was more pronounced in males, with HFD-fed β*Ahr*^WT^ males showing a significant ~1.7-fold increase in plasma insulin throughout the ipGTT relative to LFD-fed β*Ahr*^WT^ males, while HFD-fed β*Ahr*^KO^ males exhibited similar plasma insulin levels to LFD-fed β*Ahr*^KO^ males **(Figure 6Ei–iv)**. The absence of HFD-induced hyperinsulinemia in β*Ahr*^KO^ mice suggests β-cell-specific AhR signaling is required for this early compensatory response.

**Figure 6.**
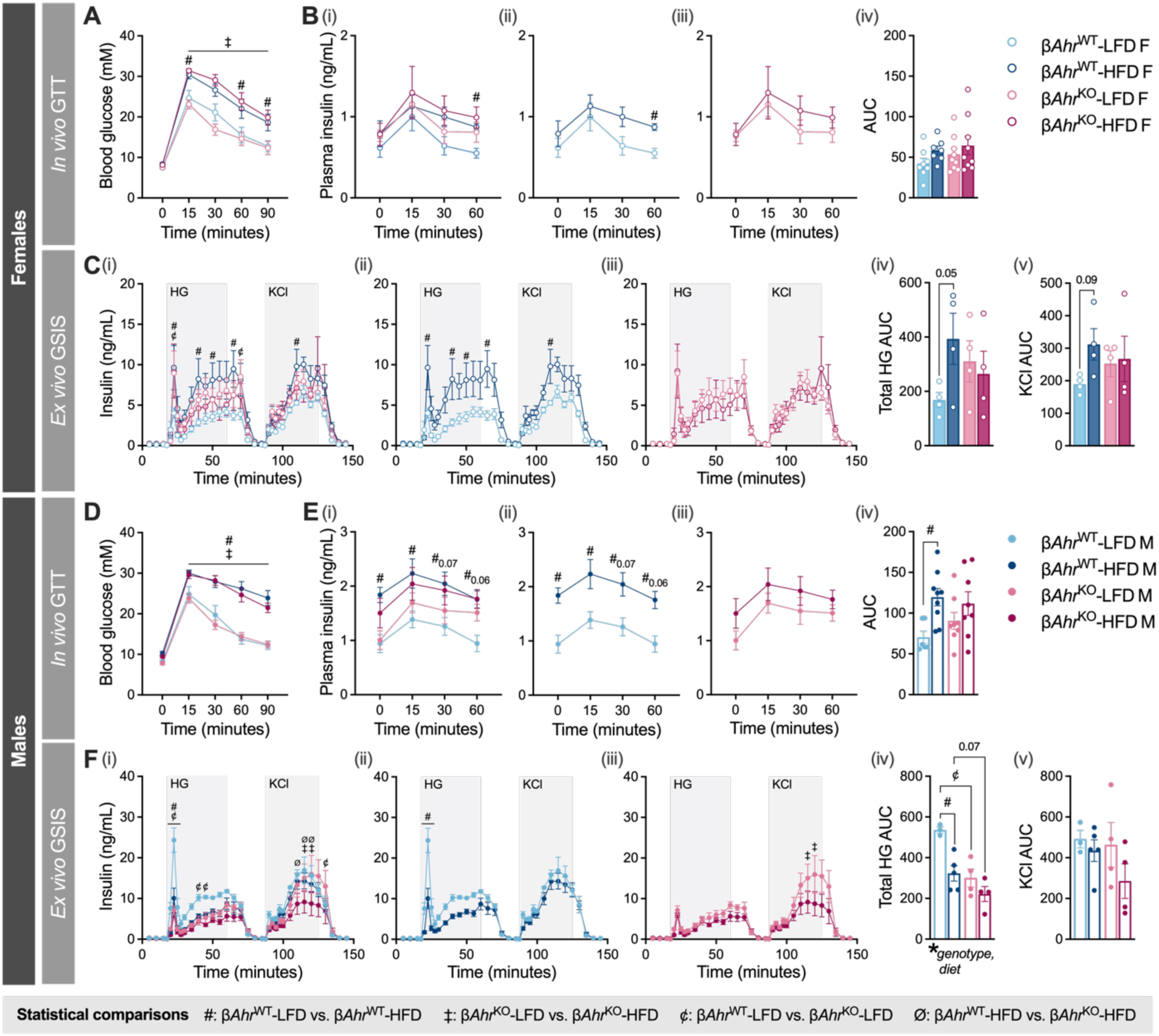
Knocking-out *Ahr* in β-cells prevented high-fat diet (HFD)-induced hyperinsulinemia in both sexes *in vivo*, but had sex-specific effects on insulin secretion *ex vivo*. A glucose tolerance test (GTT) was performed on *day 7* (*A*–*B*, *D*–*E*) and islets were isolated on *day 9* for *ex vivo* perifusion assays (*C*, *F*) in female (*A*–*C*) and male mice (*D*–*F*). Mice received a 2 g/kg bolus of glucose by intraperitoneal injection during the GTT (*n* = 7–9 per group) (*A*–*B*, *D*–*E*). Blood glucose (*A*, *D*) and plasma insulin (*B*, *E*) concentrations were measured at 15-minute intervals. For perifusion assays (*n* = 3–5 per group), insulin concentration was measured every 2.5 to 5 minutes as isolated islets were sequentially exposed to 2.8 mM low glucose, 16.7 mM high glucose (HG), or 30 mM KCl solutions (*C*, *F*). Insulin secretion profiles are shown with all groups together (*Ci*, *Fi*) or separated by genotype (*Cii*–*iii*, *Fii*–*iii*). Area under the curve (AUC) values were calculated for total HG insulin secretion (t = 17.5–80 minutes; *Civ*, *Fiv*) and KCl insulin secretion (t = 87.5–140 minutes; *Cv*, *Fv*). All data are presented as means ± SEM. Line graphs show glucose (*A*, *D*) or insulin concentrations (*B*–*C* and *E*–*F*, *i*–*iii*) over time. Line graphs for plasma insulin and *ex vivo* insulin secretion are presented with all groups combined (*B*–*C* and *E*–*F*, *i*) or separated by genotype (*B*–*C* and *E*–*F*, *ii*–*iii*). Bar graphs show AUC values, with individual data points representing biological replicates (*Biv*, *Civ*–*v*, *Eiv*, *Fiv*–*v*). The following statistical tests were used — line graphs (*A*, *Bi*–*iii*, *Ci*–*iii*, *D*, *Ei*–*iii*, *Fi*–*iii*): linear mixed-effects model followed by Tukey’s post hoc tests on estimated marginal means for pairwise comparisons at each timepoint. Bar graphs (*Biv*, *Cii*–*v*, *Eiv*, *Fii*–*v*): ordinary two-way ANOVA with uncorrected Fisher’s LSD. Statistically significant results (*p* < 0.05) are denoted by an asterisk (*) for overall two-way ANOVA effects or by the following symbols for pairwise comparisons — #β*Ahr*^WT^-LFD vs. β*Ahr*^WT^-HFD; ‡β*Ahr*^KO^-LFD vs. β*Ahr*^KO^-HFD, ¢β*Ahr*^WT^-LFD vs. β*Ahr*^KO^-HFD, Øβ*Ahr*^WT^-HFD vs. β*Ahr*^KO^-HFD.

In line with the *in vivo* plasma insulin data, HFD-induced changes in insulin secretion were only observed in β*Ahr*^WT^ islets, not in β*Ahr*^KO^ islets. Perifusion analysis using islets isolated on day 8 showed that, relative to LFD-fed controls, HFD feeding increased total HG AUC by ~2.3-fold in female β*Ahr*^WT^ islets **(Figure 6Civ)** but decreased total HG AUC by ~50% in male β*Ahr*^WT^ islets **(Figure 6Fiv)**. In contrast, HFD feeding had no effect on *ex vivo* insulin secretion in β*Ahr*^KO^ islets from either sex **(Figure 6C, F)**.

### 3.8 β*Ahr*^KO^ islets have altered expression of genes involved in the oxidative stress response compared to β*Ahr*^WT^ islets, irrespective of diet and sex

To determine whether β-cell AhR is involved in any HFD-driven transcriptional changes in islets, we performed qPCR on islets isolated on day 8 post-HFD. We measured the expression of AhR pathway genes – *Cyp1a1, Ahrr, Nrf2* **(Figure 7Ai–iii, Ci–iii)**, oxidative stress response genes – *Nqo1, Gpx1, Il-1β* **(Figure 7Aiv–vi, Civ–vi)**, β-cell identity genes – *MafA, Ins1* **(Figure 7Bi–ii, Di–ii)**, insulin processing genes – *Pcsk1, Pcsk2* **(Figure 7Biii–iv, Diii–iv)**, and glucose transport and metabolism genes – *Glut2, Gck, G6pc2* **(Figure 7Bvi–vii, Dvi–vii)**.

**Figure 7.**
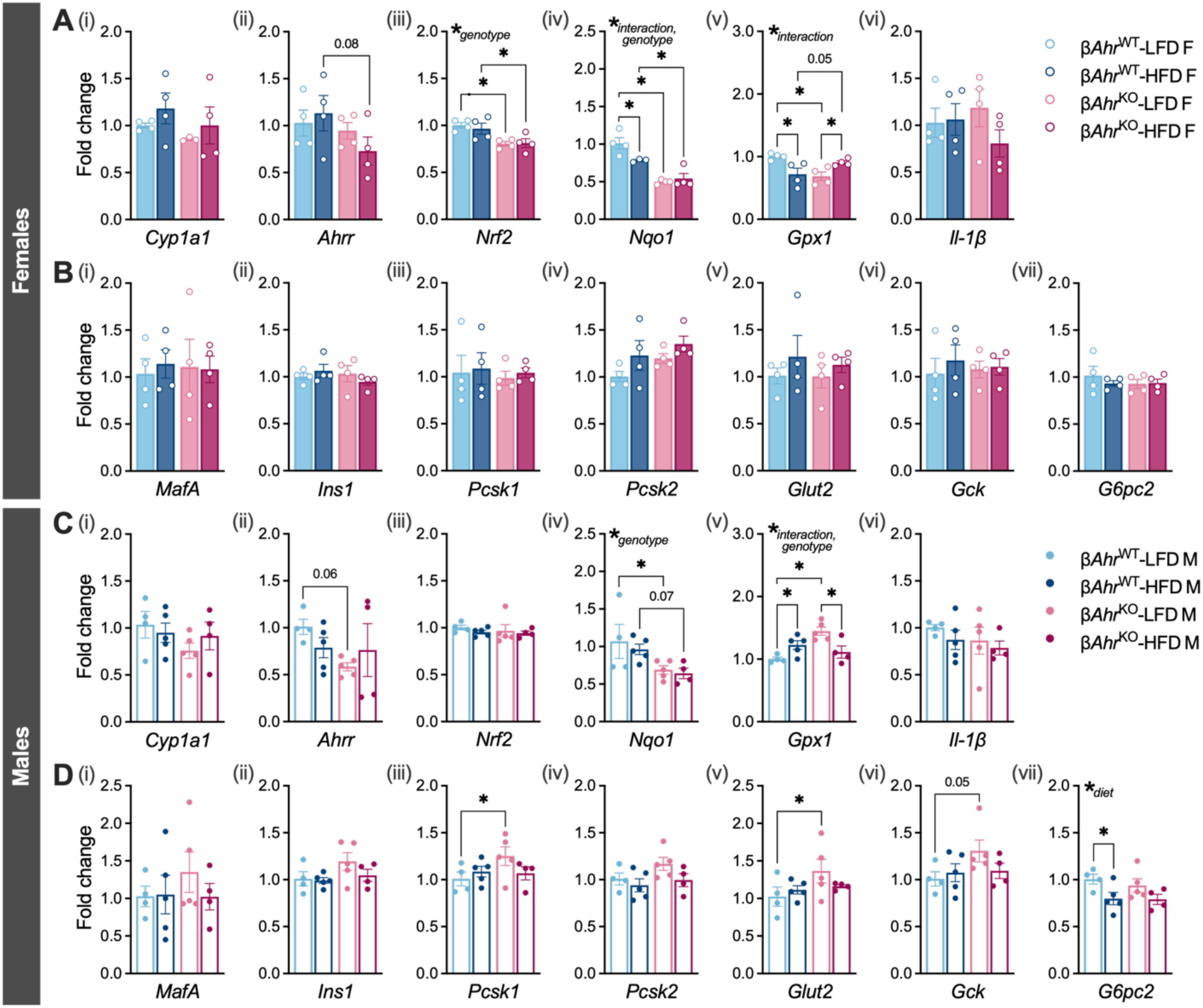
Both male and female β*Ahr*^KO^ islets have altered expression of genes involved in the oxidative stress response (*Nqo1*, *Gpx1*) compared to β*Ahr*^WT^ islets irrespective of diet. Gene expression measured by qPCR in female (*A*, *B*) and male (*C*, *D*) mouse islets (*n*_females_ = 3–4 per group, *n*_males_ = 4–5 per group). Genes involved in the AhR pathway (*A* and *C*, *i*–*iv*), oxidative stress response (*A* and *C*, *iv*–*v*), and inflammation (*A* and *C*, *i*–*vi*). Markers of β-cell maturity *(B* and *D*, *i*–*ii*) and genes involved in insulin processing (*B* and *D*, *iii*–*iv*), glucose transport (*B* and *D*, *v*), and glucose metabolism (*B* and *D*, *vi*–*vii*). All data are presented as means ± SEM; individual data points represent biological replicates. Statistical significance was assessed using an ordinary two-way ANOVA with uncorrected Fisher’s LSD; \**p* < 0.05. LFD = low-fat diet, HFD = high-fat diet.

In female islets, AhR pathway and oxidative response genes were altered by diet or genotype **(Figure 7A–B)**. In β*Ahr*^WT^ islets, HFD feeding decreased *Nqo1* and *Gpx1* expression by ~22% and ~28%, respectively. In contrast, HFD feeding had no effect on *Nqo1* expression but increased *Gpx1* expression by ~37% in β*Ahr*^KO^ islets. Under LFD-fed conditions, β-cell *Ahr* deletion reduced the expression of *Nrf2* (~20%), *Nqo1* (~50%), and *Gpx1* (~30%) **(Figure 7Aiii–v)**. In islets from HFD-fed females, β-cell *Ahr* deletion similarly reduced *Nrf2* (~16%) and *Nqo1* (~30%) expression but caused a trending increase in *Gpx1* expression (~26%) and a trending decrease in *Ahrr* expression (~36%) **(Figure 7Aii–v)**.

While gene expression changes in female islets were limited to AhR pathway and oxidative stress-related genes, we also observed changes in insulin processing and glucose metabolism genes in male islets **(Figure 7C–D)**. HFD feeding increased *Gpx1* expression (~22%) and reduced *G6pc2* expression (~21%) in β*Ahr*^WT^ islets, whereas in β*Ahr*^KO^ islets, HFD reduced *Gpx1* expression (~23%) and did not affect *G6pc2* expression **(Figure 7Cv, Dvii)**. Meanwhile, most of the genotype-driven differences were observed under LFD-fed conditions. In islets from LFD-fed males, β-cell *Ahr* deletion reduced *Nqo1* expression (~36%; **Figure 7Civ**) and increased expression of *Gpx1* (~45%; **Figure 7Cv**), *Pcsk1* (~24%; **Figure 7Diii**), and *Glut2* (~33%; **Figure 7Dv**). Additionally, islets from LFD-fed β*Ahr*^KO^ mice showed a trending decrease in *Ahrr* (~43%; **Figure 7Cii**) and a trending increase in *Gck* (~30%; **Figure 7Dvi**) relative to islets from LFD-fed β*Ahr*^WT^ mice.

### 3.9 β-cell-specific *Ahr* deletion increased alpha cell area and reduced β-cell area per islet in a diet- and sex-dependent manner

On day 8, whole pancreas was collected from a different subset of mice to assess islet size, relative α- and β-cell composition, and β-cell proliferation. Average islet area was similar across all groups in both sexes **(Figure 8A, D)**. Interestingly, LFD-fed β*Ahr*^KO^ female mice had increased % Gcg^+^ area per islet compared to LFD-fed β*Ahr*^WT^ female mice, but this difference was not observed in islets from HFD-fed mice **(Figure 8Bii)**. HFD feeding increased % Ins^+^ area per islet and decreased % Gcg^+^ area per islet in male β*Ahr*^WT^ mice, but not β*Ahr*^KO^ mice **(Figure 8Ei–ii)**. Lastly, the % Ins^+^PCNA^+^ cells was similar across all groups, indicating that β-cell proliferation was not influenced by diet or genotype in either sex at this timepoint **(Figure 8C, F, H)**.

**Figure 8.**
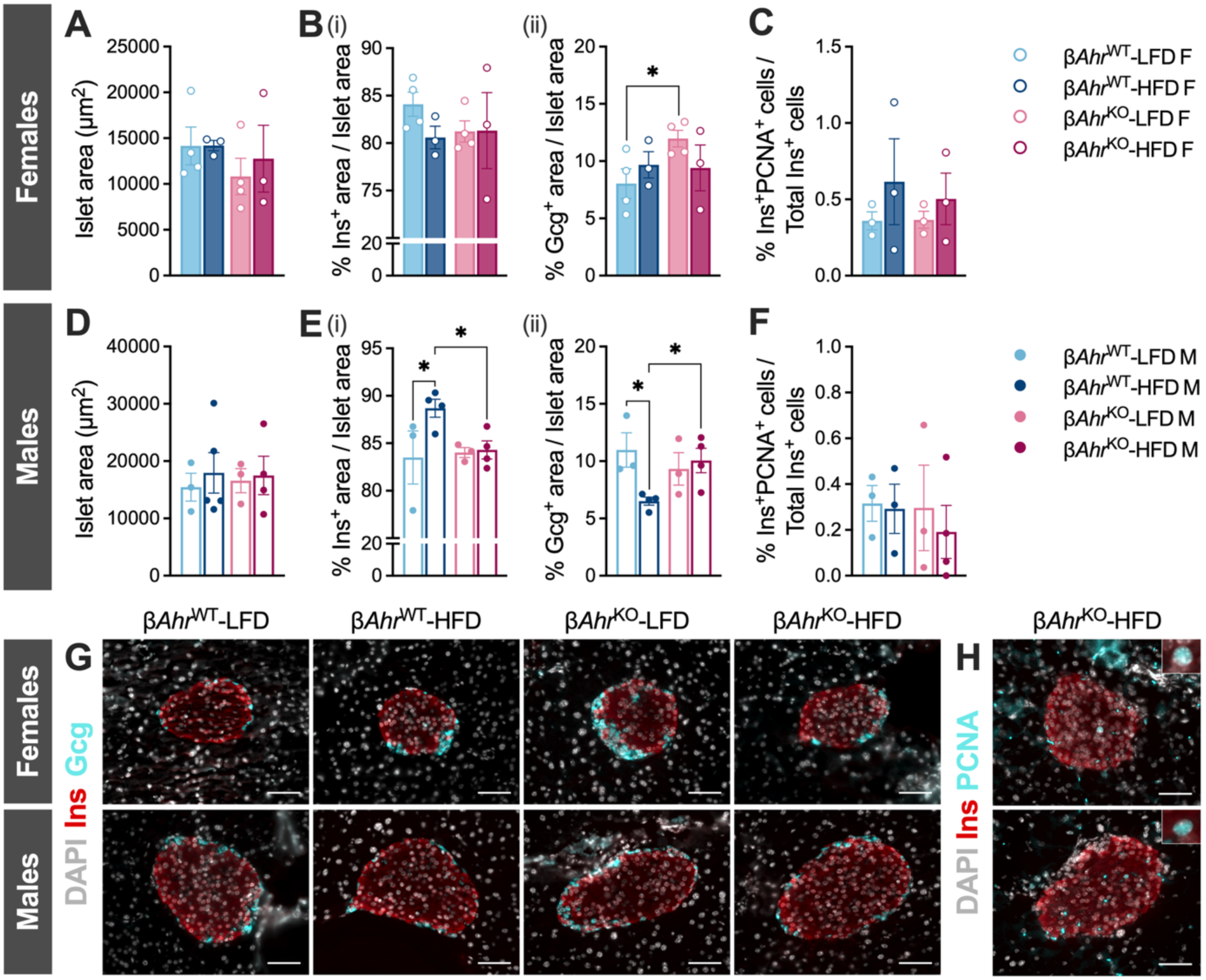
β-cell-specific *Ahr* deletion increased alpha cell area and reduced β-cell area per islet in a diet- and sex-dependent manner. Whole pancreas samples were collected on *day 9* for immunofluorescence staining. Pancreas sections from female and male mice were either co-stained with anti-insulin and anti-glucagon antibodies to assess islet morphology (*A*–*B*, *D*–*E*, *G*), or with anti-insulin and anti-PCNA antibodies to assess β-cell proliferation (*C, F, H*) (*n* = 3–4 per group). *A*–*F*: Average islet area (*A*, *D*), proportion of insulin positive (Ins^+^) area per total islet area (*Bi*, *Ei*), proportion of glucagon positive (Gcg^+^) area per total islet area (*Bii*, *Eii*), and proportion of cells positive for both insulin and PCNA (Ins^+^PCNA^+^) to the total number of cells positive for insulin (Ins^+^PCNA^+^ + Ins^+^PCNA^-^) per islet (*C*, *F*). Representative images of islets showing Ins^+^ (red) and Gcg^+^ (cyan) cells (*G*), and islets showing Ins^+^ (red) and PCNA^+^ (cyan) cells (*H*) and at 40x magnification (scale bars = 50 µM). All data are presented as means ± SEM. Individual data points represent biological replicates. LFD, low-fat diet; HFD, high-fat diet; PCNA: proliferating cell nuclear antigen. Statistical significance was determined using an ordinary two-way ANOVA with uncorrected Fisher’s LSD; \**p* < 0.05.

## DISCUSSION

We used a range of approaches—from human islet transcriptomics to *in vitro* and *in vivo* mouse experiments—to investigate the role of canonical and non-canonical AhR signaling in islet cells. Bulk human islet RNA-seq revealed enrichment of pathways involved in xenobiotic and nutrient metabolism in individuals with higher *CYP1A1* expression, suggesting that the AhR–CYP1A axis may influence broader metabolic processes beyond its role in xenobiotic detoxification. *In vitro* experiments showed global *Cyp1a1/1a2* deletion increased the susceptibility of female islets to xenobiotic stress but had minimal effect on the responses of islets to glucolipotoxicity in either sex. In contrast, β-cell *Ahr* deletion altered islet responses to glucolipotoxic stress but did not affect xenobiotic stress responses. Furthermore, β-cell *Ahr* deletion prevented the compensatory hyperinsulinemia observed in HFD-fed wildtype mice of both sexes *in vivo*, although the effect was more pronounced in males. Our data suggest that AhR signaling contributes to islet biology in a context- and sex-dependent manner.

Our results demonstrate that endogenous AhR pathway activity in human islets is associated not only with xenobiotic metabolism, but also with pathways involved in nutrient metabolism. Interestingly, xenobiotic metabolism pathways were enriched exclusively among upregulated genes, whereas nutrient metabolism pathways were enriched among both up- and downregulated genes. This pattern suggests that the AhR–CYP1A axis is associated with coordinated induction of detoxification genes but more complex regulation of nutrient metabolism in human islets. Among the most enriched pathways in upregulated genes was retinol metabolism, which has been implicated in metabolic diseases, including obesity and T2D [37,38]. Furthermore, *ALDH1A3*, one of the upregulated genes from the retinol metabolism pathway in *CYP1A1-*high donors, has been associated with β-cell dedifferentiation and reduced insulin secretion [39,40]. Another notable gene was *G6PC1*, which is part of the glycolysis/gluconeogenesis pathway. *G6PC1* is not normally expressed in islets and is typically restricted to liver, kidneys, and intestines [41]; therefore, its upregulation in donors with high *CYP1A1* may reflect loss of β-cell identity or metabolic reprogramming. Lastly, while exploratory, our sex-stratified analyses suggest that CYP1A1-mediated AhR activity is more closely associated with hormone and nutrient metabolism in females, and with xenobiotic metabolism and immune-related pathways in males. Altogether, these findings support a role for the AhR–CYP1A axis in integrating detoxification and nutrient-related processes in human islets, potentially with sex-specific effects. While promising, further investigation is necessary to validate the involvement of AhR signaling in retinol metabolism, nutrient processing, and maintaining β-cell identity in human islets.

Our lab previously showed that *in vitro* TCDD exposure impairs GSIS in *Cyp*^WT^ but not *Cyp*^KO^ male islets [14], suggesting that CYP1A enzymes mediate the deleterious effects of TCDD on islet function. We expanded on this work by including both female and male mice and by using perifusion analysis instead of static GSIS to better characterize insulin secretion dynamics. Contrary to our previous findings, TCDD did not impair insulin secretion in either *Cyp*^WT^ or *Cyp*^KO^ male islets, suggesting that the effect of TCDD on GSIS in males may be modest and sensitive to cohort-level variation so it is difficult to detect consistently across experiments. Interestingly, TCDD modestly impaired 1^st^ phase GSIS in female *Cyp*^WT^ islets, a phenotype that was worsened in *Cyp*^KO^ islets. Furthermore, TCDD modestly increased maximal mitochondrial respiration and spare respiratory capacity in *Cyp*^WT^ female islets, but not in *Cyp*^KO^ female islets. Together, these results suggest that *Cyp1a1/1a2* expression in female islets is protective under chemical conditions, potentially by maintaining mitochondrial adaptability. Since the genotype effect in females was most evident during the 2^nd^ phase of GSIS and in response to KCl, future studies could examine whether altered calcium dynamics, granule recruitment, or other downstream exocytotic mechanisms contribute to this phenotype.

Exposure to high-glucose, high-fatty acid (i.e., GLT) media elicited similar functional responses in *Cyp*^WT^ and *Cyp*^KO^ islets, irrespective of sex. These findings suggest that the AhR–CYP1A axis does not play a major role in islet responses to acute glucolipotoxicity for 48 hours *in vitro*. However, when *Cyp*^WT^ and *Cyp*^KO^ mice were fed 45% HFD for 14 weeks, *Cyp*^KO^ males exhibited delayed HFD-induced hyperinsulinemia. Moreover, HFD feeding impaired GSIS in *Cyp*^WT^, but not *Cyp*^KO^, male islets [29]. The incongruence in these findings might reflect technical and temporal differences between acute (48 hour) *in vitro* nutrient stress and chronic (14 week) *in vivo* metabolic challenge, as well as systemic effects driven by the global *Cyp*^KO^ in other tissues. Variations in nutrient composition between rodent diets and culture media could also be playing a role, given that islets can respond differently to specific nutrients [42,43]. Follow-up experiments could investigate the involvement of the AhR–CYP1A axis in the islet response to other types of nutrients (e.g., amino acids) and nutrient mixtures. Overall, our results point to a nuanced role for the AhR–CYP1A axis in islet biology, likely supporting long-term metabolic adaptation rather than directly mediating acute responses to glucolipotoxicity.

We previously showed that a single high-dose TCDD injection alters glycemia in female and male β*Ahr*^WT^ mice and impairs insulin secretion in isolated male β*Ahr*^WT^ islets [21]. Importantly, these TCDD-induced phenotypes were absent in β*Ahr*^KO^ mice, indicating that β-cell *Ahr* mediates these effects [21]. In comparison, 48 hours of *in vitro* TCDD treatment did not alter insulin secretion in β*Ahr*^WT^ or β*Ahr*^KO^ islets of either sex. This suggests that the direct effects of TCDD on islets may be modest and difficult to detect, or that systemic factors that are missed with an *in vitro* model contribute to TCDD-induced islet dysfunction. The absence of a TCDD-driven phenotype *in vitro* also limits our ability to determine whether β-cell *Ahr* is involved in islet responses to direct TCDD exposure. Together, these findings underscore the importance of using both *in vivo* and *in vitro* models to distinguish context-dependent responses from robust phenotypes.

Our data highlight the importance of β-cell *Ahr* in basal islet function and nutrient stress responses, with sex-specific effects. In male islets, β-cell *Ahr* appears to protect against GLT-induced dysfunction, whereas in female islets, β-cell *Ahr* deletion consistently increased basal insulin secretion across exposure models. This phenotype in β*Ahr*^KO^ female islets aligns with previous findings from our group [21] and with reports of increased insulin secretion following loss of AhR in INS1 cells with an *AHR* knockdown and in human stem cell-derived islets with a global *AHR* knockout [44,45]. Following HFD, β-cell *Ahr* deletion prevented adaptive changes in plasma insulin and islet insulin secretion in both sexes. While β*Ahr*^WT^ mice developed HFD-induced compensatory hyperinsulinemia *in vivo*, this adaptation was not observed in β*Ahr*^KO^ mice. Likewise, islets isolated from HFD-fed β*Ahr*^WT^ mice exhibited sex-specific changes in insulin secretion that were absent in β*Ahr*^KO^ islets. These genotype-driven differences in plasma insulin and islet function coincided with modest transcriptional changes in the oxidative stress–related genes, which may have contributed to the impaired capacity of β*Ahr*^KO^ islets to meet the increased metabolic demand of HFD feeding. Overall, our results demonstrate that β-cell *Ahr* is critical for mounting adaptive hyperinsulinemia in both sexes. We also show that β-cell AhR signaling has sex-specific roles, primarily supporting nutrient stress adaptation males, while also regulating basal insulin secretion in females.

In addition to its role in regulating insulin secretion, β-cell *Ahr* may also influence islet remodelling. In males, HFD-feeding increased the proportion of β-cells and decreased the proportion of α-cells per islet in β*Ahr*^WT^ mice, but did not alter islet cell proportions in β*Ahr*^KO^ mice. These changes in male islet composition parallel the *in vivo* phenotype, in which HFD-induced hyperinsulinemia was observed in β*Ahr*^WT^ mice but not β*Ahr*^KO^ mice, further supporting a role for β-cell *Ahr* in early adaptation to HFD. Because male mice lacking β-cell *Ahr* failed to exhibit hyperinsulinemia or an increase in β-cell area following HFD, we hypothesized that β-cell *Ahr* deletion may prevent adaptive β-cell proliferation. Although we did not observe any changes in β-cell proliferation on day 9 post-HFD, we may have missed the peak window of proliferation, which has been reported to occur as early as 3 days after HFD feeding [46]. In females, diet did not affect islet cell proportions; instead, β-cell *Ahr* deletion increased the proportion of α-cells per islet only under LFD-fed conditions. Since glucagon can potentiate insulin secretion [47], this increase in relative α-cell area may partly explain the enhanced insulin secretion observed in β*Ahr*^KO^ female islets. Together, our findings suggest that β-cell *Ahr* contributes to adaptive islet remodelling in males and to baseline islet physiology in females. To more comprehensively assess the role of β-cell *Ahr* in proliferation, future studies can use S961, an insulin receptor agonist that robustly induces β-cell proliferation, together with *in vivo* labeling using a modified thymidine analog to capture proliferative events over time instead of at a single timepoint [48,49].

By examining both canonical and non-canonical AhR signaling using complementary model systems, our study provides novel insight into the role of AhR signaling in normal islet physiology and stress responses. While the canonical AhR–CYP1A axis primarily mediates the xenobiotic stress response, broader AhR signaling in β-cells also regulates responses to nutrient stress. Our results also reveal sex-specific effects. For instance, the AhR–CYP1A axis may play a greater role in supporting resilience to chemical stress in female islets than in male islets. Moreover, in females, β-cell *Ahr* supports both baseline islet physiology and nutrient stress responses, whereas in males its role is largely limited to the latter. Overall, our findings identify AhR signaling as a regulator of islet responses to chemical and metabolic stress. Because humans are routinely exposed to both metabolic stressors and environmental AhR ligands, determining how AhR signaling integrates these exposures to shape metabolic responses will be important for clarifying its contribution to T2D risk.

## 4. ACKNOWLEDGEMENTS

We extend our sincere gratitude to all the donors and their families for their invaluable contribution to diabetes research. We thank the Lund University Diabetes Centre, supported by the Nordic Network for Clinical Islet Transplantation, for generating and making available the bulk human islet RNA-seq dataset deposited in GEO under accession GSE50398. We also thank the HumanIslets.com Consortium for providing human bulk islet RNA-seq data from a Canadian cohort. HumanIslets.com was funded by the Canadian Institutes of Health Research, JDRF Canada, and Diabetes Canada (5-SRA-2021-1149-S-B/TG 179092), with data generated from islets isolated by the ADI IsletCore with support from the Human Organ Procurement and Exchange (HOPE) program, Trillium Gift of Life Network (TGLN), and other Canadian organ procurement organizations. Donor consent was obtained in writing, as approved by the Human Research Ethics Board at the University of Alberta (Pro00013094).

We also thank the University of Ottawa Bioinformatics Core and the Stem Cell Network for providing critical training and support in RNA-seq analysis. In particular, we are grateful to Gareth Palidwor for retrieving the primary RNA-seq dataset from GEO and providing the processed count data, and to Christopher Porter for sharing his RNA-seq analysis pipeline, which we adapted for this study. Finally, we thank Dr. Sanjeena Dang, Associate Professor and Canada Research Chair at Carleton University, for sharing her expertise in statistics and bioinformatics to help us finalize our analyses.

## 7. AUTHOR CONTRIBUTIONS

M.E.A.C., M.P.H., and J.E.B. conceived and designed research; M.E.A.C., M.P.H., L.B., J.P., L.G., R.T., A.K., and E.P.-G. performed experiments; M.E.A.C., M.P.H., and J.E.B. analyzed data; M.E.A.C., M.P.H., and J.E.B. interpreted results of experiments; M.E.A.C. prepared figures; M.E.A.C. and J.E.B. drafted manuscript; M.E.A.C., M.P.H., L.B., J.P., L.G., E.v.Z., and J.E.B. edited and revised manuscript; M.E.A.C., M.P.H., L.B., J.P., L.G., R.T., A.K., E.P.-G. and J.E.B. approved final version of manuscript.

