## Supplemental materials for "β-cell-specific *Ahr* expression is critical to high-fat diet-induced hyperinsulinemia"

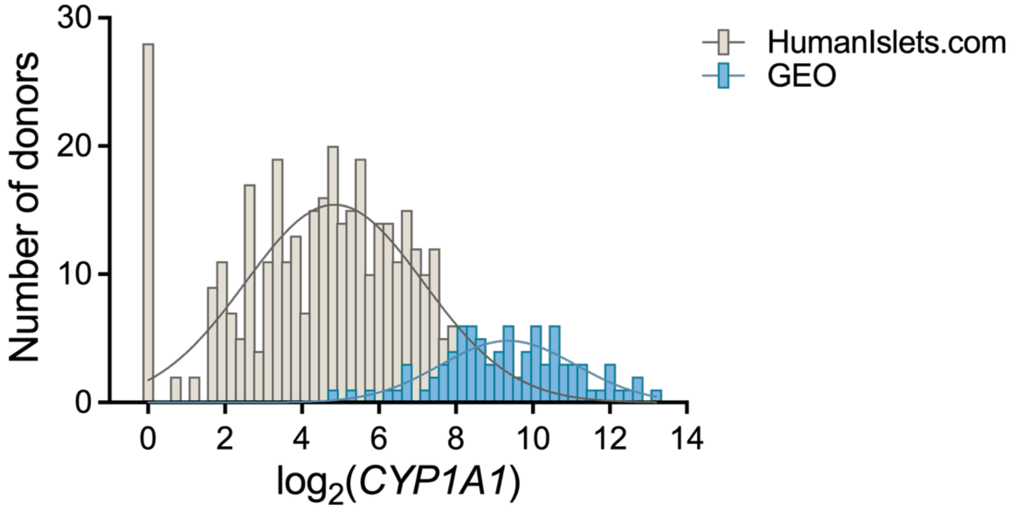


**Supplementary Figure 1. *CYP1A1* expression is generally higher in donors from the Gene Expression Omnibus (GEO) dataset compared with donors in the HumanIslets.com dataset.** Gene counts were normalized using *DESeq2* (v1.46.0) and $\log_2$-transformed as $\log_2$(*CYP1A1* + 1).

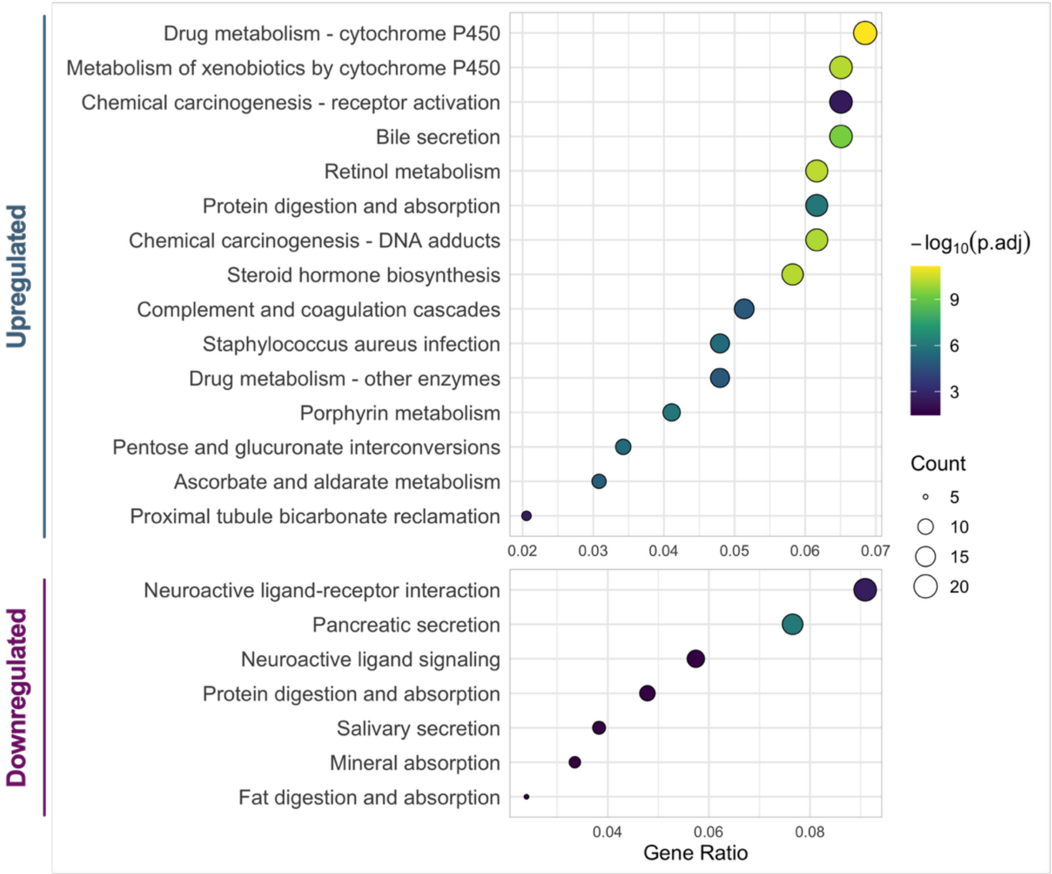


**Supplementary Figure 2. Pathway enrichment analysis results after excluding the middle 20% of donors (n = 18) based on $\log_2$(*CYP1A1*) expression distribution.** Only donors in the top and bottom 40% were included in the analysis (n = 35 per group). The dot plot represent pathways enriched among upregulated differentially expressed genes (DEGs) (top) or downregulated DEGs (bottom), ordered by increasing GeneRatio. DEGs were identified using *DESeq2* (v1.50.2), pathway enrichment was performed using *clusterProfiler* (v4.18.4), and the graphs were generated with *enrichplot* (v1.30.5). All analyses were performed in R (v4.5.3) using RStudio (v2026.04.0+526).

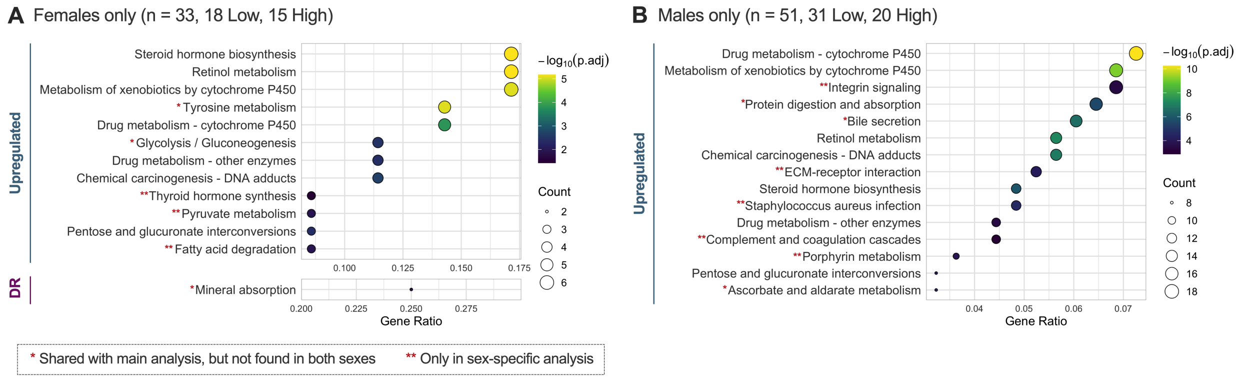


**Supplementary Figure 3. Sex-stratified pathway enrichment analysis.** After excluding 4 donors with intermediate *CYP1A1* levels, 33 females and 51 males remained. Within each sex, donors were divided into "low" and "high" *CYP1A1* groups based on $\log_2$(*CYP1A1*) expression. Differential gene expression and over-representation analyses were performed separately for females (A) and males (B). Dot plots represent pathways enriched among upregulated differentially expressed genes (DEGs) or downregulated DEGs (DR), ordered by increasing GeneRatio. DEGs were identified using *DESeq2* (v1.46.0), pathway enrichment was performed using *clusterProfiler* (v4.14.6), and graphs were generated with *enrichplot* (v1.26.6). All analyses were performed in R (v4.4.2) using RStudio (v2025.05.0+496).

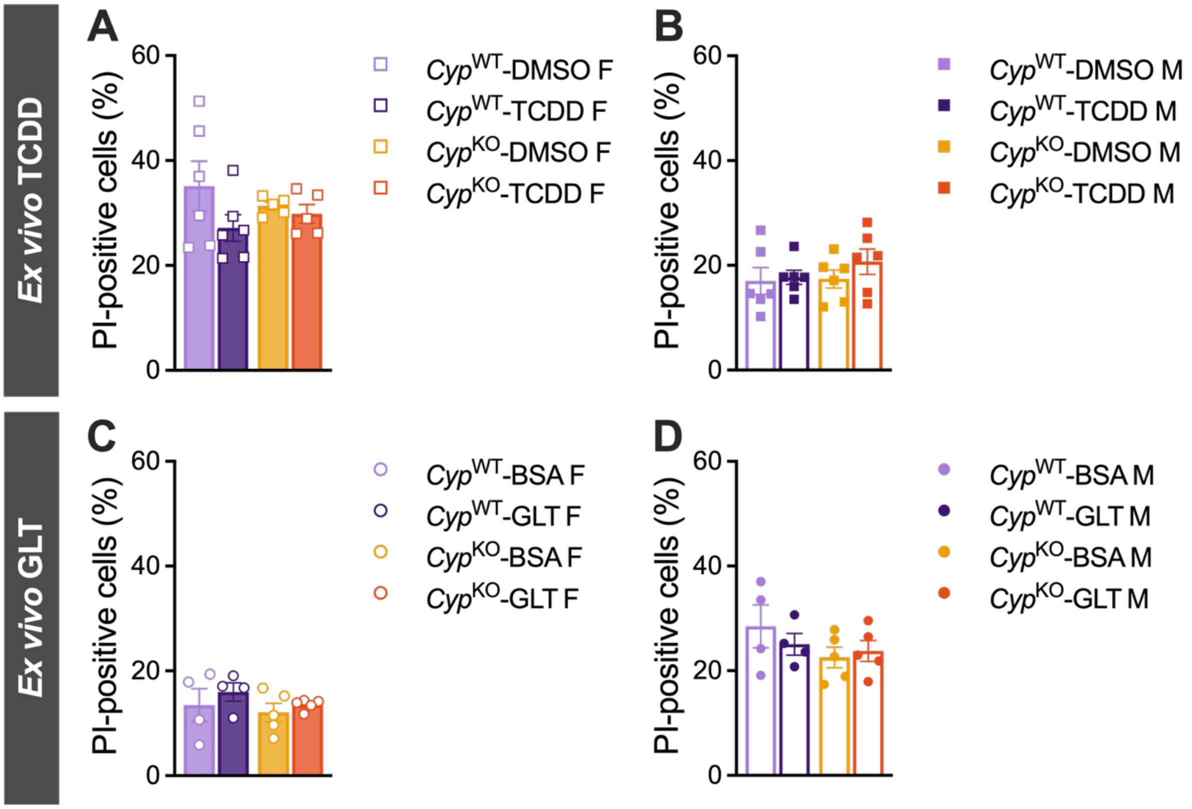


**Supplementary Figure 4. Islet cell viability was not affected by *ex vivo* treatment with chemical or glucolipotoxicity (GLT).** After 48h of *ex vivo* 2,3,7,8-tetrachlorodibenzo-*p*-dioxin (TCDD) (*A* and *B*) or GLT exposure (*C* and *D*), $Cyp^{WT}$ and $Cyp^{KO}$ islets were dispersed into a single cell suspension to assess cell viability with propidium iodide (PI)-inclusion assays. A microscopy-based PI-inclusion assay was used for TCDD experiments (*A* and *B*; n = 5–6 per group), and a flow-cytometry-based PI-inclusion assay was used for GLT experiments (*C* and *D*; n = 4–5 per group). Images from the microscopy-based PI-inclusion assay were quantified semi-automatically by a blinded single observer using the Zeiss Zen Blue 2.6 software. Statistical test used: repeated measures two-way ANOVA with uncorrected Fisher's LSD at $p < 0.05$. DMSO = dimethyl sulfoxide, BSA = bovine serum albumin.

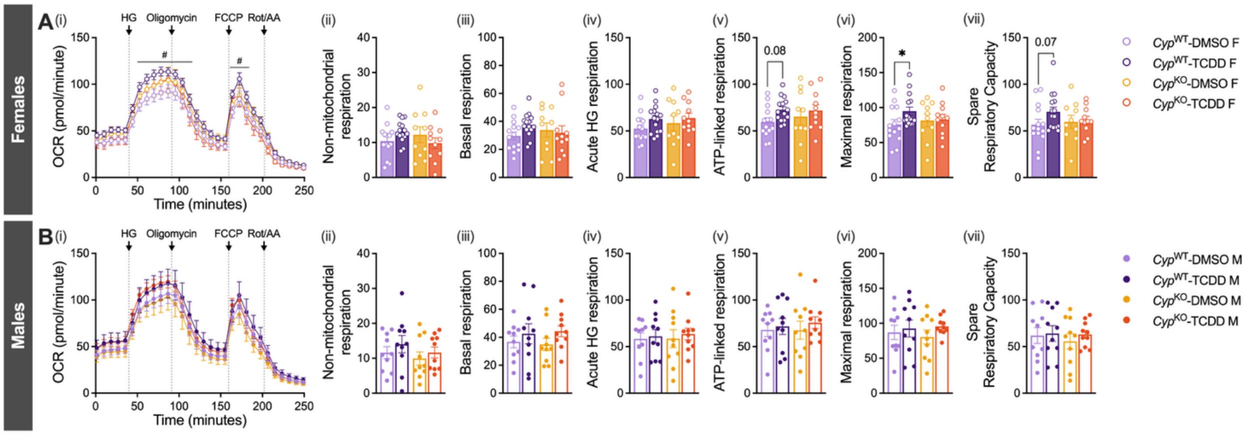


**Supplementary Figure 5. TCDD exposure increased maximal respiration in *Cyp*$^{\text{WT}}$ female islets but not in *Cyp*$^{\text{KO}}$ female islets.** Mitochondrial stress tests were performed on *Cyp*$^{\text{KO}}$ and *Cyp*$^{\text{WT}}$ islets ($n_{\text{females}}$ = 10–14 per group, $n_{\text{males}}$ = 10 per group) using the Agilent Seahorse XFe24 Analyzer (*A*, *D*). Oxygen consumption rate (OCR) was measured as islets were subjected to sequential injections of HG solution, oligomycin, FCCP, and rotenone/antimycin (Rot/AA) (*i*). Non-mitochondrial respiration (*ii*), basal respiration (*iii*), acute HG respiration (*iv*), ATP-linked respiration (*v*), maximal respiration (*vi*), and spare respiratory capacity (*vii*) were calculated using formulas provided by Agilent. The following statistical tests were used — line graphs (*i*): linear mixed-effects model followed by Tukey's post hoc tests on estimated marginal means for pairwise comparisons at each timepoint; bar graphs (*ii*–*vi*): mixed-effects model with uncorrected Fisher's LSD. All data are presented as means ± SEM. Individual data points on bar graphs (*ii*–*vii*) represent biological replicates.

**Supplementary Table 1. Top 50 differentially expressed genes (DEGs) between "low *CYP1A1*" and "high *CYP1A1*" donors that were mapped to KEGG pathways during over-representation analysis.** $Log_2FC$ = $log_2FoldChange$, $p_{adj}$ = FDR-adjusted p-value.

| Gene Symbol | Gene Name | $log_2FC$ | $p_{adj}$ |
|---|---|---|---|
| *ADH1A* | alcohol dehydrogenase 1A (class I), alpha polypeptide | 2.98 | 1.14E-07 |
| *KRT20* | keratin 20 | 2.93 | 9.74E-09 |
| *CYP1A1* | cytochrome P450 family 1 subfamily A member 1 | 2.90 | 2.00E-41 |
| *SI* | sucrase-isomaltase | 2.84 | 1.71E-03 |
| *UGT1A1* | UDP glucuronosyltransferase family 1 member A1 | 2.75 | 8.20E-05 |
| *CYP17A1* | cytochrome P450 family 17 subfamily A member 1 | 2.64 | 8.00E-04 |
| *KCNJ13* | potassium inwardly rectifying channel subfamily J member 13 | 2.29 | 3.46E-06 |
| *CYP2B6* | cytochrome P450 family 2 subfamily B member 6 | 2.17 | 2.73E-05 |
| *MYO1A* | myosin IA | 2.16 | 3.59E-07 |
| *CYP4F2* | cytochrome P450 family 4 subfamily F member 2 | 2.08 | 2.37E-04 |
| *CREB3L3* | cAMP responsive element binding protein 3 like 3 | 2.07 | 5.45E-04 |
| *OTC* | ornithine transcarbamylase | 2.06 | 3.43E-06 |
| *ADH1C* | alcohol dehydrogenase 1C (class I), gamma polypeptide | 1.88 | 1.04E-06 |
| *AKR1C4* | aldo-keto reductase family 1 member C4 | 1.85 | 2.29E-03 |
| *GUCA2A* | guanylate cyclase activator 2A | 1.82 | 1.68E-05 |
| *UGT2B17* | UDP glucuronosyltransferase family 2 member B17 | 1.80 | 6.30E-03 |
| *IYD* | iodotyrosine deiodinase | 1.79 | 2.87E-05 |
| *CASP5* | caspase 5 | 1.79 | 6.07E-03 |
| *UGT1A8* | UDP glucuronosyltransferase family 1 member A8 | 1.79 | 2.46E-04 |
| *C8A* | complement C8 alpha chain | 1.76 | 2.24E-02 |
| *G6PC1* | glucose-6-phosphatase catalytic subunit 1 | 1.76 | 1.14E-07 |
| *DMBT1* | deleted in malignant brain tumors 1 | 1.75 | 1.03E-04 |
| *AQP1* | aquaporin 1 (Colton blood group) | 1.62 | 9.91E-06 |
| *KIF20A* | kinesin family member 20A | 1.53 | 3.60E-03 |
| *UGT2B11* | UDP glucuronosyltransferase family 2 member B11 | 1.52 | 4.99E-02 |
| *ADH6* | alcohol dehydrogenase 6 (class V) | 1.52 | 9.82E-07 |
| *VSIG4* | V-set and immunoglobulin domain containing 4 | 1.50 | 1.24E-03 |
| *UGT2B15* | UDP glucuronosyltransferase family 2 member B15 | 1.49 | 1.26E-05 |
| *ALDH3A1* | aldehyde dehydrogenase 3 family member A1 | 1.46 | 3.05E-03 |
| *FGF19* | fibroblast growth factor 19 | 1.45 | 1.21E-03 |
| *PLA2G2A* | phospholipase A2 group IIA | 1.43 | 4.21E-02 |
| *HSD17B2* | hydroxysteroid 17-beta dehydrogenase 2 | 1.41 | 3.07E-07 |
| *ST6GALNAC1* | ST6 N-acetylgalactosaminide alpha-2,6-sialyltransferase 1 | 1.37 | 2.90E-04 |
| *ALB* | albumin | 1.36 | 1.13E-02 |
| *CDH17* | cadherin 17 | 1.36 | 1.03E-05 |
| *SFRP4* | secreted frizzled related protein 4 | -1.83 | 5.47E-04 |
| *MT1H* | metallothionein 1H | -1.74 | 7.09E-03 |
| *CELA3B* | chymotrypsin like elastase 3B | -1.69 | 2.22E-03 |
| *P2RX1* | purinergic receptor P2X 1 | -1.65 | 8.16E-04 |
| *MT1G* | metallothionein 1G | -1.65 | 8.60E-05 |
| *RBPJL* | recombination signal binding protein for immunoglobulin kappa J region like | -1.59 | 1.29E-02 |
| *CLDN18* | claudin 18 | -1.59 | 1.88E-03 |
| *IGF1* | insulin like growth factor 1 | -1.53 | 3.64E-02 |
| *C7* | complement C7 | -1.51 | 6.11E-04 |
| *CXCL14* | C-X-C motif chemokine ligand 14 | -1.51 | 9.74E-03 |
| *HTR2A* | 5-hydroxytryptamine receptor 2A | -1.51 | 3.20E-04 |
| *ATP4A* | ATPase H+/K+ transporting subunit alpha | -1.49 | 6.34E-03 |
| *GAL* | galanin and GMAP prepropeptide | -1.48 | 4.43E-05 |
| *GSTT2* | glutathione S-transferase theta 2 (gene/pseudogene) | -1.46 | 2.11E-02 |
| *ARHGDIG* | Rho GDP dissociation inhibitor gamma | -1.36 | 7.52E-03 |

**Supplementary Table 2.** Equations used to calculate mitochondrial function assay parameters. The formulas were adapted from the Report Generator User Guide for the Agilent Seahorse Cell Mito Stress Test. OCR = oxygen consumption rate, Rot/AA = rotenone and antimycin A, HG = high glucose, FCCP = carbonyl cyanide-4 (trifluoromethoxy) phenylhydrazone.

| Parameter | Equation |
|---|---|
| Non-mitochondrial respiration | Minimum OCR after Rot/AA injection |
| Basal respiration | (Last OCR measurement before HG injection) – (Non-mitochondrial oxygen consumption) |
| Acute HG respiration | (Last OCR measurement before Oligomycin injection) – (Last OCR measurement before HG injection) |
| ATP-linked respiration | (Last OCR measurement before Oligomycin injection) – (Minimum OCR measurement after Oligomycin injection) |
| Maximal respiration | (Maximum OCR measurement after FCCP injection) – (Non-mitochondrial oxygen consumption) |
| Spare respiratory capacity | (Maximal respiration) – (Basal respiration) |

**Supplementary Table 3.** Primer sequences used for qPCR.

| Gene | Forward sequence | Reverse sequence | Amplicon size (bp) |
|---|---|---|---|
| *Ahrr* | GGTGGGTCGCTGAGTATGGT | CGGTTGCTCTTACCGGAAGC | 162 |
| *Cyp1a1* | ATCACAGACAGCCTCATTGAGC | AGATAGCAGTTGTGACTGTGTC | 139 |
| *G6pc2* | AGCTGCCCTAAGCTACACCA | AACACTCCACAGAAAGGACCA | 85 |
| *Gck* | TGGTGGATGAGAGCTCAGTG | TGAGCAGCACAAGTCGTACC | 96 |
| *Glut2* | GCAACTGGGTCTGCAATTTT | CCAGCGAAGAGGAAGAACAC | 88 |
| *Gpx1* | CTCACCCGCTCTTTACCTTC | CACACCGGAGACCAAATGATG | 103 |
| *Il-1β* | GCCACCTTTTGACAGTGATGAG | AGCTTCTCCACAGCCACAAT | 186 |
| *Ins1* | TCAGAGACCATCAGCAAGCA | CTCCCAGAGGGCAAGCAG | 89 |
| *MafA* | AGTCGTGCCGCTTCAAG | CGCCAACTTCTCGTATTTCTCC | 149 |
| *Nqo1* | CTCTGGCCGATTCAGAGTGG | GTCTCCTCCCAGACGGTTTC | 152 |
| *Nrf2* | CTCTGCTGCAAGTAGCCTCG | TGTCTTGCCTCCAAAGGATGT | 138 |
| *Pcsk1* | GGTGGAAAGGTCGAGTCTAGC | TGCACACCAAACGCAAAAGA | 153 |
| *Pcsk2* | TTTGGAGTCCGAAAGCTCCC | GGTGTAGGCTGCGTCTTCTT | 91 |
| *Ppia* | GCCAGGACCTGTATGCTTTA | AGCTCTGAGCACTGGAGAGA | 178 |
